# Convergence and novelty in adaptation to whole genome duplication in three independent polyploids

**DOI:** 10.1101/2020.03.31.017939

**Authors:** Sian M. Bray, Eva M. Wolf, Min Zhou, Silvia Busoms, Magdalena Bohutínská, Pirita Paajanen, Patrick Monnahan, Jordan Koch, Sina Fischer, Marcus A. Koch, Levi Yant

## Abstract

Convergent evolution is observed broadly across the web of life, but the degree of evolutionary constraint during adaptation of core intracellular processes is not known. High constraint has been assumed for conserved processes, such as cell division and DNA repair, but reports of nimble evolutionary shifts in these processes have confounded this expectation. Whole genome duplication (WGD) necessitates the concerted adjustment of a wide range of fundamental intracellular functions but nevertheless has been repeatedly survived in all kingdoms. Given this repeated adaptation to WGD despite obvious intracellular challenges to core processes such as meiosis, we asked: how do lineages not only survive WGD, but sometimes ultimately thrive? Are the solutions employed constrained or diverse? Here we detect genes and processes under selection following WGD in the *Cochlearia* species complex by performing a scan for selective sweeps following WGD in a large-scale survey of 73 resequenced individuals from 23 populations across Europe. We then contrast our results from two independent WGDs in *Arabidopsis arenosa* and *Cardamine amara.* We find that while WGD does require the adaptation of particular functional processes in all three cases, the specific genes recruited to respond are highly flexible. We also observe evidence of varying degrees of convergence between different cases. Our results point to a polygenic basis for the distributed adaptive systems that control meiotic crossover number, ionomic rewiring, cell cycle control, and nuclear regulation. Given the sheer number of loci under selection post-WGD, we surmise that this polygenicity may explain the general lack of convergence between these species that are ~30 million years diverged. Based on our results, we speculate that adaptive processes themselves – such as the rate of generation of structural genomic variants—may be altered by WGD in nascent autopolyploids, contributing to the occasionally spectacular adaptability of autopolyploids observed across kingdoms.

---

Biologists have long been fascinated by the convergent evolution of similar traits in distant lineages^1^. Given the tractability of population-based resequencing studies to detect candidate mechanisms underlying adaptations, the genomic underpinnings of convergence are increasingly coming to light^2^. Diverse examples can be found in all kingdoms, from the convergent basis for the loss in flight in birds^3^, to the evolution of toxin-resistant herbivory in insects^4^, to drug resistance in pathogens^5^. These studies ask the fundamental question: to what degree is evolution constrained along a given adaptive trajectory? That is, do convergent traits, when detected, have a basis in similar or identical genetic changes? Ultimately, this drives at a bigger question: is evolution predictable? Overwhelmingly, these works focus on adaptation to external selection pressures. Here, in contrast, we investigate convergence in genomic responses to an array of internal physiological pressures resulting from whole genome duplication (WGD).

The duplication of an entire genome is a dramatic mutation that disrupts the most fundamental of cellular processes, yet is full of promise for those that can adapt to a transformed WGD state^6–8^. Immediately following WGD in autopolyploids (within species WGD, with no hybridisation), a series of novel challenges arise. The most obvious concerns meiosis: the instant doubling of chromosome homologs complicates their neat pairing during meiosis^9^, with consequences that directly reduce fitness. If a chromosome engages in crossovers with more than one other homolog, the likelihood of entanglement dramatically rises, along with breakage upon anaphase. Simultaneously, WGD throws evolved cellular equilibria off balance, including ion homeostasis, protein expression and cell size regulation^6,7,10,11^. These challenges are so severe that they are insurmountable for many young autopolyploids, although established polyploid populations persist in nature for many species, indicating that the early challenges can be overcome.

To date there have been two genome-scale investigations probing the genomic basis of adaptation to WGD. The first evidence for specific adaptative signatures in response to WGD comes from *Arabidopsis arenosa^12,13^.* Population resequencing studies scanning for divergent selection across the genomes of this young autotetraploid revealed a set of physically and functionally interacting proteins exhibiting the strongest signatures of selection post-WGD. While a range of processes was under selection, 8 of the 18 most robust signatures of selective sweep directly overlapped genes that appear to have coevolved to decrease chromosome crossover rates during prophase I of meiosis^13^. This same suite of alleles was found to be shared in a sister species, *Arabidopsis lyrata*, with which *A. arenosa* hybridises in the wild, specifically between the autotetraploid cytotypes of each species^14^. The importance of these genes was highlighted by the discovery that these same alleles were shared by both young autotetraploids. Specific signatures of gene flow at these same 8 alleles indicated that the two cases were not independent, and that these 8 alleles that cooperatively function had their origins in sperate diploid species, coming together across species barriers only when the two autotetraploids hybridised^15,16^. More recently, a pool-seq-based genome scan for divergence outliers following WGD in *Cardamine amara*, a Brassicaceae ~17 million years diverged from *A. arenosa*, was unable to detect excess convergence beyond that expected by chance among individual loci under selection at the gene orthologue level^17^. However, a modest degree of convergence on the level of functional pathways was detected, in particular for genes that control meiosis. Despite this convergence at the process level, the marked coevolution of functionally and physically interacting chromosome crossover-governing genes discovered in *A. arenosa* was largely absent in *C. amara^17^.*

Here we test whether this convergence signal is further abrogated in a more distantly related, independent WGD event. With the addition of a third case we can better estimate whether adaptation of this set of interacting meiosis proteins is the exception or the rule. We focus here on the *Cochlearia* species complex, which is ~20-25 million years diverged from *A. arenosa*^18–21^. The *Cochlearia* genus exhibits two ploidy series with diploid base chromosome number n=6 and n=7, with the n=6 series consisting of diploid, tetraploid, hexaploid, octoploid and eventually heptaploid cytotypes, which broadly hybridise in nature^22–28^. This cytotype richness is magnified by the presence of B chromosomes in some populations^29^. *Cochlearia* is found across Europe, from Spain to the Arctic, in a wide range of habitats including freshwater springs, coastal cliffs, sand dunes, salt marshes, metal contaminated sites and roadside grit^22,24,33–38,26–31,31,32^. A broad habitat differentiation is evident by ploidy, with diploids typically found in upland freshwater springs, tetraploids on coasts, often directly adjacent to seawater or continuously submerged, and hexaploids in similarly extreme saline conditions. In fact, the hexaploid *C. danica* is one of the most rapidly spreading invasive species in the UK and Continental Europe, specifically invading salted roadways since the 1970’s^39,40^, and thriving the most highly saline road grit conditions^41,42^.

We first assess *Cochlearia* demography and scan for selective sweeps post-WGD by individually resequencing 76 individuals from 23 diploid, tetraploid, and hexaploid populations sampled from across Europe, focusing analysis on diploids and tetraploids in the UK. We find evidence of convergence at the functional level across all three species and partial convergence at the gene ortholog level between *Cochlearia* and *A. arenosa*, but not *C. amara*. This suggests that the same core set of cellular functions must adapt in response to WGD, but that the specific genes that can be utilized to this end are not fully constrained, though there is some evidence of ortholog-level convergence.

## Results and Discussion

### Population sampling and genome assembly

To determine optimal populations to contrast for WGD-specific signatures of selection, we first sampled populations across the reported range of the *Cochlearia* species complex throughout Europe^24,26,41,43,44,27,28,30,31,31–34^ and then conducted a flow cytometry-based survey of genome size variation (Dataset S1). Measurements were normalised against the diploid population with the most stable individual within-population genome size estimates, WOL. Because *Cochlearia* species extensively hybridise and exhibit considerable phenotypic plasticity, we here primarily designate populations by demonstrated ploidy rather than taxonomic names: in general diploids = *Cochlearia pyrenaica*, tetraploids = *Cochlearia officinalis*, hexaploids = *Cochlearia danica* (coastal dunes and roadside) or octoploids = *Cochlearia anglica* (immediately coastal, marshes).

We constructed a *de novo* genome assembly of one diploid individual from the NEN population using 10x Genomics Chromium linked-read sequencing assembled with Supernova 2.0.0 (91% complete BUSCOs; contig n50=40kbp; Table S1, Table S2 and Methods). We next choose for population-level genome resequencing 116 individuals from 25 populations across Europe and re-sequenced these using Illumina PE format (genome-wide average sequencing depth = 15x; Figure 1A and Table S3). After retaining only individuals with a minimum average genome-wide sequencing depth (average=21x; min=4x), this cohort consisted of 76 individuals from 23 populations. The final dataset consisted of 6,020,948 SNPs (quality and depth filtered; see Methods).

**Fig. 1.**
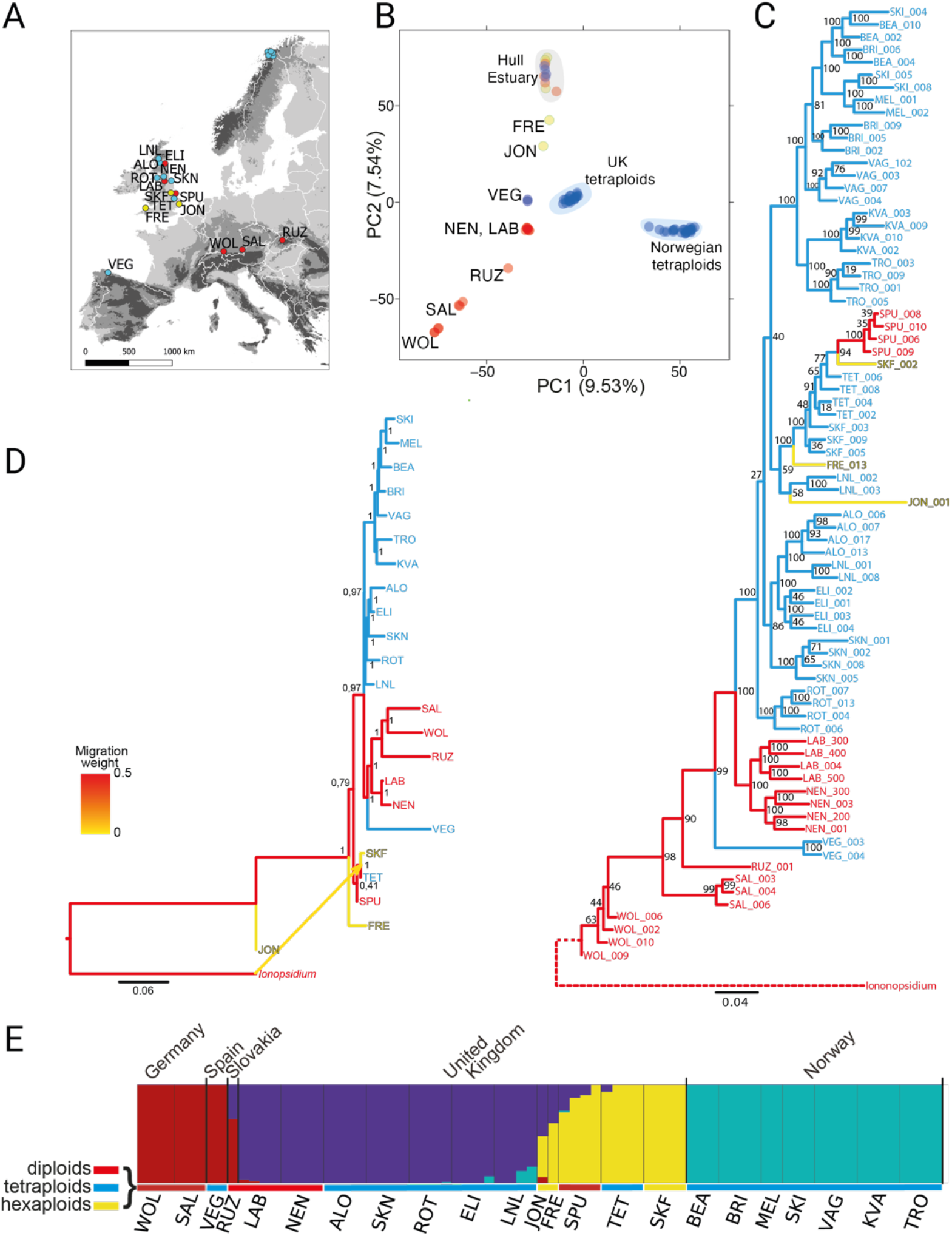
Geographic distribution and genetic structure of *Cochlearia.* **A,** distribution of the 23 *Cochlearia* populations (red labels, diploids; blue, tetraploids; yellow, hexaploids; locations are given in Table S3). **B,** Principal Component Analysis (PCA) of all populations. **C,** Rooted phylogeny of 76 individuals from 23 populations created with RAxML, with outgroup *Ionopsidium.* **D,** TreeMix graph of populations, indicating migration edge from *Ionopsidium* to SKF/TET ancestor. **E,** fastSTRUCTURE analysis (k=4, min alleles=8) of all *Cochlearia* individuals, with regions, populations, and ploidies indicated.

### Demographic structure and WGD selection scan population choice

To assess demographic structure, we first performed principal component analysis (PCA) on 415,139 (putatively neutral) 4-fold-degenerate SNPs. The first two axes indicate that geographic origin dominates over ploidy: PC1 (9.5% of variability) differentiates populations by geographic location, and PC2 (7.5% of variability) differentiates by ploidy (figure 1B). The PC1/PC2 distribution resembles a three-pointed star (concave hexagon), where each point represents one ploidy over a gradient of geographic distance, with the exception of individuals from the Hull estuary on the east of Northern England, where individuals of all ploidies intermingle, suggesting extensive local interploidy introgression and a complex reticulate system. Global ancestry estimation with fastSTRUCTURE was consistent with this observation (Fig. 1E), showing obvious clustering by ploidy and geographic location as well as interploidy admixture, especially in the Hull region.

Next, we performed a phylogenetic analysis in RAxML based on 72,641 4-fold-degenerate SNPs (using the relative *Ionopsidium* to root the tree; Figure 1C). Bootstrap values were strong for major groupings (e.g. the clustering of British diploids vs. tetraploids or the Norwegian tetraploids), although backbone resolution was weaker, with the positioning of groups flipped between trees. Such a pattern could result from high levels of introgression or rapid radiation. Introgression seems likely, as there is known interploidy hybridisation in *Cochlearia*^23–25,28^. This observation is supported by the admixture seen by fastSTRUCTURE and SplitsTree analyses (Figure 1E and 2C). This could also be consistent with a rapid inter- or peri-glacial radiation^18^ and postglacial migration and introgression scenarios such as found in other *Cochlearia* species^28^. To further assess demographic history and admixture patterns, we performed TreeMix modeling (on 52,186 biallelic, fourfold-degenerate SNPs), which represents genome relationships through a graph of ancestral populations^45^. Here, the optimal number of migration edges was determined to be a single one (based on the Evanno method) which revealed an admixture signal from the outgroup *Ionopsidium* to British *Cochlearia* hexaploids SKF (Figure 1D). Taken together, these demographic analyses indicate complex patterns of ancestry and/or hybridization among the hexaploids, but simpler groupings between diploids and tetraploids.

**Fig. 2.**
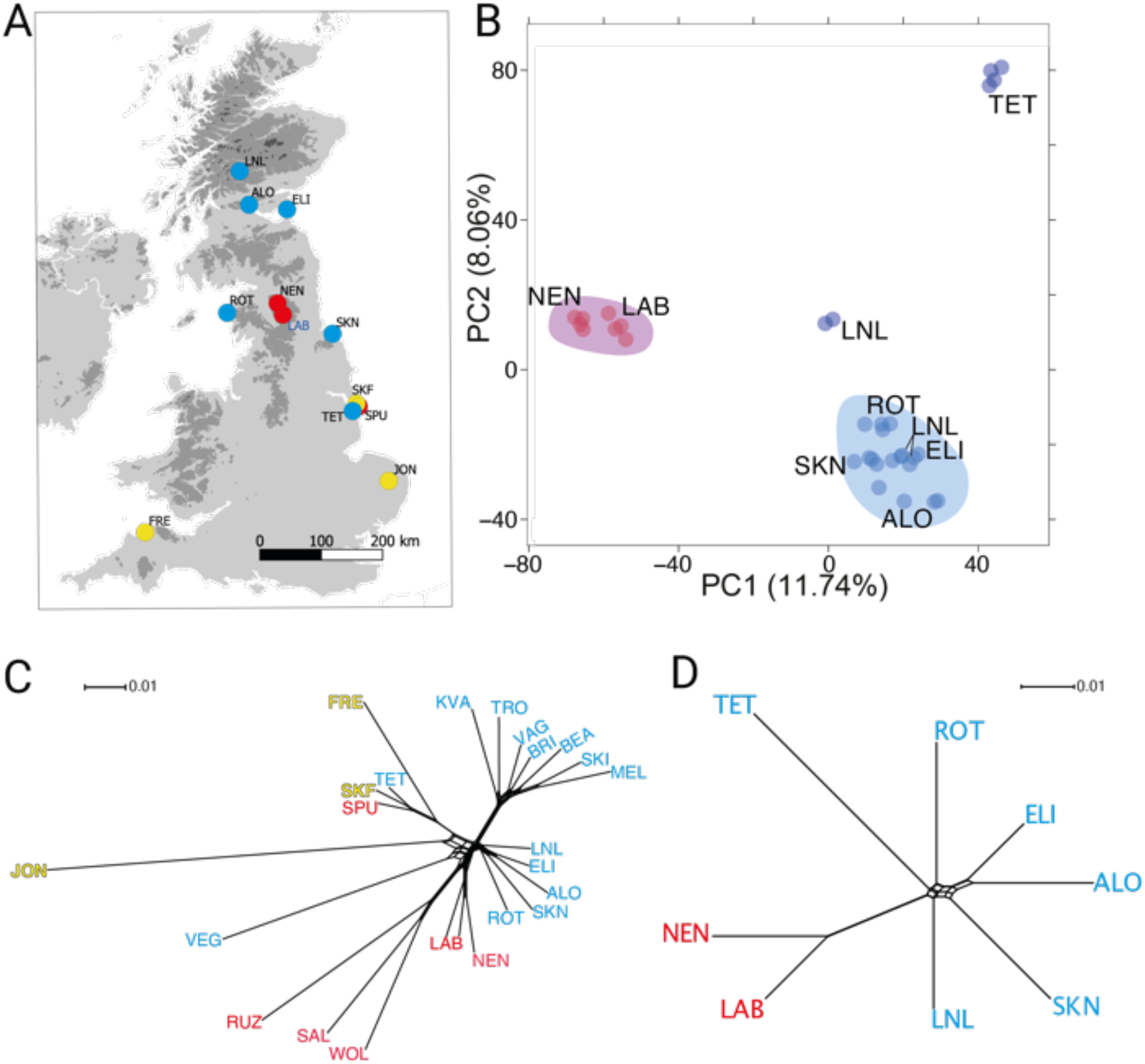
British *Cochlearia* demographic structure. **A,** British *Cochlearia* populations used for selective sweep scan (red labels, diploids; blue, tetraploids; yellow, hexaploids). **B,** PCA of British samples only, excluding hexaploids. **C** and **D,** Nei’s genetic distances visualised in a SplitsTree representation for all sequenced populations (C), and only British diploids and tetraploids (B).

To focus on adaptation following WGD, we selected six sequenced populations of British *Cochlearia* tetraploids (27 individuals; average depth >20x) and two populations of British *Cochlearia* diploids (8 individuals; average depth >20x; Fig. 2A). This resulted in a dataset containing 6,020,948 SNPs after quality and depth filtering. To assess population relationships in this simplified demographic scenario, we first performed a PCA using 361,981 4-fold-degenerate SNPs. The first axis (11.7% of variability explained) clearly separated the samples by ploidy, while the second (8.1% of variability explained) separated the LNL and TET individuals that showed signs of admixture with either the Norwegian tetraploids or British hexaploids respectively (Fig. 2B-D, Fig. 1E).

### Selective sweeps associated specifically with WGD

To determine which genes exhibit the strongest signatures of selection following WGD in *Cochlearia* tetraploids, we contrasted allele frequencies across the genomes of our British diploids and tetraploids, calculating differentiation metrics (Rho^46^, Hudson’s Fst^47^, Nei’s Fst^48^, Weir-Cochran’s Fst^49^, Dxy^50^, number of fixed differences and average groupwise allele frequency difference) for all 1 kb windows genome-wide which contain a minimum of 20 post-filter SNPs, a minimum average depth of 8x and a maximum of 20% missing data. The number of SNPs in this contrast was 3,024,896 residing in 44,968 windows, covering 39,594 of the 44,023 predicted genes, or 90% of all gene coding loci.

To determine which differentiation metric most reliably identified genomic regions that exhibit peaks in allele frequency difference (AFD) above local background levels, we performed a quantitative inspection of all AFD plots in the outlier tails of empirical distributions of each differentiation metric (see Methods). Based on superior performance in this assessment, we used Hudson’s Fst^47^, which brings the added benefit of robustness for unequal population sizes and presence of rare variants^51^. Given that Fst-based selective sweep scans have met with success in other diploid/autopolyploid systems^12,13,15,17,52^, this also makes our current results directly comparable to previous works. We extracted windows in the top 1% of the Fst distribution as empirical outliers, consisting of 450 1kb windows, overlapping 296 gene-coding loci (Dataset S2). This list was further refined using a fineMAV-like method^17,53^, which uses Grantham scores to predict the potential functional impact of each SNP that encodes a non-synonymous amino acid change. This approach then amplifies the severity of each predicted amino acid change by the AFD between the SNPs encoding the change. Out of the 448,625 non-synonymous-encoding SNPs assigned a MAV score, the 1% outliers from the empirical distribution were reserved (4,486 SNPs; Dataset S3) and intersected with our 296 Fst outlier windows, yielding a refined list of 144 gene coding loci, containing 406 MAV SNPs (bold in Dataset S2). A selection of AFD peaks for these candidate genes exhibiting gene-localised, ploidy-specific selective sweep signatures is given in Fig. 3.

**Fig. 3.**
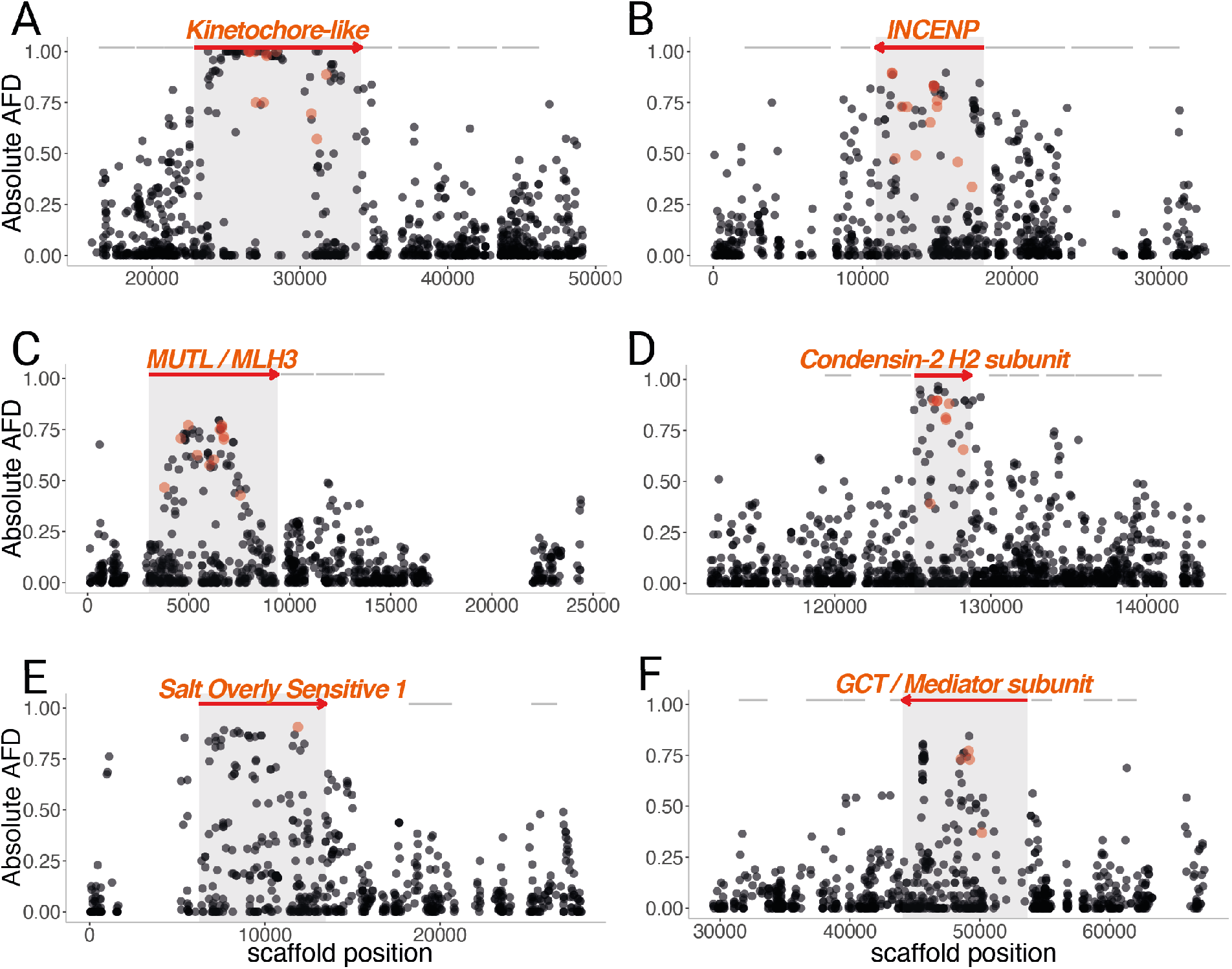
Selective sweep signatures at DNA management and ion homeostasis loci. Examples of selective sweep signatures among 6 Fst outlier genes (red arrows). The X axis gives genome position in base pairs. The Y axis gives AFD values at single SNPs (dots) between diploid and tetraploid *Cochlearia.* Red arrows indicate genes overlapping top 1 % Fst windows and grey lines indicate neighboring genes. Light red dots indicate MAV outlier SNPs present in the candidate selected locus.

### Functional processes under selection post-WGD in *Cochlearia*

Using our 296 WGD-specific selective sweep candidates, gene ontology (GO) enrichment analysis yielded 100 significantly enriched biological processes (using a conservative ‘elim’ Exact Test p < 0.05, Table S4; see Methods). Taking higher level categories (minimum 150 annotated genes/category) yielded 30 significantly enriched biological processes (Table S4, bold rows). Among the most significantly enriched categories we observe DNA integration, cell division, meiotic and sister chromatid segregation, mitosis, DNA repair, and recombination represented. Other processes such as response to salt stress, organelle fission, stomatal complex development, and cellular cation homeostasis were also significantly enriched. Overall, these processes are remarkably well aligned with physiological changes post-WGD suspected to be important challenges to the establishment of nascent autopolyploids^8,11^.

### Evolution of DNA movement and related processes

Eleven of the top sixteen most enriched higher-level GO categories were terms closely related to cell division, organelle fission, cell cycle and DNA metabolic processes, including DNA recombination and DNA repair, and cellular response to DNA repair (Table S4). Candidate genes contributing to these enrichments have predicted roles in cell division, specifically meiosis, or chromosome movement more generally. For example, a particularly dramatic selective sweep signature overlaps directly *CPg20426* (the orthologue of *AT3G10180)* and includes 12 MAV outlier SNPs, 6 of which are fixed in the tetraploid relative to the diploid (Fig. 3A). This gene encodes a kinetochore-like protein, which consists of a kinesin domain and a binding domain with homology to a TATA binding protein, separated by a Structural Maintenance of Chromosomes (SMC) coiled coil domain. Taken together, this suggests that the encoded protein may be part of a complex that mediates chromosome movement. The category ‘formation of the centromere’ is represented by three further genes, *CPg14121, CPg15411* and *CPg17406. CPg14121* is the ortholog of *CENP-C*, an essential kinetochore component^54,55^ and *CPg15411* is the ortholog of the inner centromere protein INCENP (Fig. 3B). Implicated in the control of chromosome segregation and cytokinesis in yeast and animals, plant INCENP has a role in the development and differentiation of the gametophytes^56–58^. Both of these loci contain 1% outlier-MAV SNPs, with the *INCENP* ortholog harboring a remarkable 14 MAV outlier SNPs, the greatest number of MAV outlier SNPs of any gene in the entire genome. Beyond these, there are a diversity of proteins with putative roles in cell division, such as CPg25053 and CPg25685 whose *A. thaliana* homologs both regulate cell cycle progression^59,60^.

### Evolution of DNA repair

Several of the most confident signals of selective sweep are found in DNA repair-related genes. In particular *CPg6863*, the ortholog of *A. thaliana MLH3* (*MutL*) is both a 1% Fst outlier and contains 12 MAV SNPs, joint third most (Fig. 3C; equal with the kinetochore protein *CPg20426* as the gene with the third highest genome-wide quantity of MAV SNPs). This mismatch repair gene controls the number of meiotic crossovers in *A. thaliana*, with mutants exhibiting a 60% reduction in crossover number^61^, making this a particularly strong candidate for modulating post-WGD meiosis, where a reduction in crossover number stands as the strongest candidate mechanism for re-establishing fertility in *A. arenosa™’^62^*.

Two other genes directly implicated in DNA repair stand out: *CPg1416*, the ortholog of the condensin-2 H2-subunit required for proper DNA double-strand break repair in *A. thaliana* and humans^63,64^, is an Fst outlier with 8 MAV SNPs (Fig. 3D). Furthermore condensin-2 has been recently implicated in controlling the association and dissociation of centromeres^65^. Additionally, we find an Fst outlier peak over CPg36347, homologue of *DAYSLEEPER*, an essential domesticated transposase^66^. *DAYSLEEPER* was first isolated as binding the Kubox1 motif upstream of the DNA repair gene *Ku70*. The complex Ku70/Ku80 regulate non-homologous end joining double-strand break repair, the only alternative to homologous recombination^66–68^.

### Selection on proteins involved in global transcription

In a polyploid the total DNA content doubles but the protein content and cell size does not scale accordingly, so we predict that the control of gene expression, like meiosis, should undergo adaptive evolution post-WGD. This is supported by empirical findings in A. arenosa^12,13,69^. Here we confirm this finding in *Cochlearia*, with 11 predicted DNA/RNA polymerase-associated genes *(CPg1405, CPg1875, CPg5061, CPg12069, CPg16591, CPg17556, CPg21554, CPg26775, CPg26954, CPg28073 and CPg31859)* and 3 putative ribosomal genes (*CPg28891, CPg34724 and CPg35322*) among our selective sweep outliers. These include *CPg1405*, the ortholog of *NRPB9A*, an RNA polymerase subunit that is implicated in transcription initiation, processivity, fidelity, proofreading and DNA repair^70–74^. We also detect an ortholog of *MED13, CPg28073*, a component of the mediator complex, which is essential for the production of nearly all cellular transcripts.

### Selection on ion transport and stress signaling

The ionomic equilibrium of the cell is immediately altered upon WGD^10^. We see signatures of selection that may represent a response to this, including stress response genes that are triggered in response to environmental ionic stressors. For example, the ortholog of *SALT OVERLY SENSITIVE 1*, a membrane Na+/H+ transporter that removes excessive Na+ from the cell^75^ and is central to salt tolerance, exhibits a selective sweep signature (Fig 3E). We also find the ortholog of *DEAD-BOX RNA HELICASE 25*, identified in *A. thaliana* as a repressor of stress signaling for salt, osmotic, and cold stress^76,77^. This gene also controls freezing tolerance^78^, a phenotype relevant to the likely cold-loving demographic history of *Cochlearia.* Similarly, we see *CPg16997*, the ortholog of drought response gene *AtHB7*^79^. To confirm that this was not the result of ecotype differences we performed a salt tolerance experiment on diploid and tetraploid plants. Surprisingly, given their divergent ecotype preferences, with tetraploids found in more saline conditions, we found that the diploid *Cochlearia* are in fact more salt tolerant than the tetraploids (p-value = 2.178e-05; See Supplementary Text 1 and Table S5.). This finding also contrasts strongly to observations of increased salinity tolerance in neotetraploid *Arabidopsis thaliana*^10^.

### Stomata, plastid-related, and other categories under selection

Several genes involved in stomatal function were outliers post-WGD, such as the ortholog of *OST2 (OPEN STOMATA2; CPg30015)*, which encodes the AHA1 protein, the major H+ ATPase in the plasma membrane that drives hyperpolarization and initiates stomatal opening. This protein is a target of ABA stress signaling to close the stomata during drought response^80^. We confirmed this functionally, detecting increases in both in stomatal conductance and net photosynthetic rate under drought conditions in tetraploid *Cochlearia* populations relative to diploids (See Supplementary Text 2, Figure S1; Tables S6-S8). Another equilibrium disrupted by WGD, and that has not been discussed in previous WGD adaptation genome scans^12,13,15,17^, is that between the chloroplast, mitochondrion and nuclear genomes. Many genes related to the function of such organelles are divergence outliers in tetraploid *Cochlearia*. For example, we detect 11 genes annotated as linked to the function of plastids/chlorophyll *(CPg1559, CPg1878, CPg2251, CPg16297, CPg18478, CPg19736, CPg21595, CPg26364, CPg30068, CPg30733* and *CPg33711)* and five linked to mitochondrial function *(CPg2266, CPg24404, CPg26437, CPg28878, CPg17406*). Notable also is that four selective sweep candidates encode myosins (*CPg10091, CPg22983, CPg6763* and *CPg35628*), suggesting a WGD associated adaptation in cellular organization, a category that encompasses organelle localization, cytoskeletal dynamics and nuclear shape^11^.

### Ortholog-level convergence

To test for convergence in adaptation to WGD at the ortholog level, we determined orthogroups using the three genomes gene annotations with OrthoFinder^81^, giving a total of 21,619 orthogroups (Methods). Top 1% Fst gene-coding outliers for *A. arenosa* (n=452; Dataset S4), *C. amara* (n=229; Dataset S5), and *Cochlearia* (n=296; Dataset S2) were extracted and then considered orthologues if they were part of the same orthogroup in the genome-wide orthofinder analysis. By this analysis, not a single orthogroup was represented in all three independent WGDs. However, there were a handful of orthogroups common to any two WGD adaptation events: 6 orthogroups were identified in both *C. amara* and *A. arenosa* outliers, while 11 were identified in both *Cochlearia* and *A. arenosa* outlier lists (Table S9). No orthogroups were common in both *Cochlearia* and *C. amara* lists. Consistent with our previous work^17^, this overlap was not significant for *C. amara* vs. *A. arenosa* (SuperExactTest P=0.23). In contrast, however, we did detect a significantly greater number of overlaps at the gene ortholog level between Fst outliers in *Cochlearia* and those in *A. arenosa* (SuperExactTest P=0.013). Gene coding loci under selection post-WGD in both *Cochlearia* and *A. arenosa* have inferred roles in DNA and RNA polymerisation (either *DNA pol V* or *nuclear DNA-directed RNA polymerase NRPB9* orthologs, respectively), potassium homeostasis (the *HIGH-AFFINITY K+ TRANSPORTER 1* ortholog), and stomatal control (the ortholog of *OPEN STOMATA 2*) (Table S9). All of these genes are involved in processes that have been implicated in adaptation to WGD^7,8,11^. These loci therefore stand as strong candidates in salient challenges to nascent polyploids. We note, however, that the degree of this convergence is low, consisting of only 3.7% of genes exhibiting the strongest signatures of selection in *Cochlearia*, and only 2.4% of those under strong selection in *A. arenosa*. While we expect that this likely represents a bona fide lack of convergence in these adaptations at the level of orthologous loci, we note that we focused here on genic signal, and there are many levels upon which selection may act beyond the scope of this study. For example, many gene regulatory changes would escape the notice of our scan, given that we required any outlier divergence window to at least partially overlap a gene coding locus^66^.

### Functional process-level convergence

While we detected minimal (or no) convergence in any of our pairwise contrasts at the gene level, we reasoned that there may be similarities in processes under selection between the three independent WGDs, representing process level, or functional convergence. To estimate this, we performed GO enrichment analysis on each outlier set and overlapped the results from each GO WGD (Table 1). Three high-level terms were significantly enriched in all three species: ‘DNA metabolic process’, ‘cellular response to abscisic acid stimulus,’ and ‘cellular response to alcohol’. Additionally, there were six subcategories that were enriched in *Cochlearia* and one of the other two species. These included DNA recombination, drought tolerance, mitosis and meiosis. Taken together, these results strongly imply that adaptive evolution in response to WGD is focused on particular functions, but that there is a high degree of stochasticity in which exact genes evolve.

**Table 1.** Enriched GO terms common among independent WGD events

|  |  | *Cochlearia* | *C. amara* | *A. arenosa* |
| --- | --- | --- | --- | --- |
| GO:0006259 | DNA metabolic process | 0.01764 | 6.50E-08 | 0.00082 |
| GO:0071215 | cellular response to ABA | 0.03738 | 0.04813 | 0.04013 |
| GO:0097306 | cellular response to alcohol | 0.03738 | 0.04813 | 0.04013 |
| GO:0051301 | cell division | 0.00044 | - | 0.0032 |
| GO:1903047 | mitotic cell cycle process | 0.00797 | - | 0.00148 |
| GO:0006310 | DNA recombination | 0.01543 | - | 0.02109 |
| GO:0000280 | nuclear division | 0.03186 | - | 4.10E-08 |
| GO:0051276 | chromosome organization | - | 0.01902 | 0.00021 |
| GO:0009738 | ABA-activated signalling | - | 0.03186 | 0.02214 |
| GO:0051321 | meiotic cell cycle | 0.0395 | 0.02571 | - |
| GO:0009414 | water deprivation response | 0.04327 | 0.04782 | - |

**Note:** GO Fisher’s Exact test elim values are given. This conservative method tests for enrichment of terms from the bottom of the GO hierarchy to the top and discards any genes that are significantly enriched in a descendant GO term.

## Conclusion

Following WGD, an instantly changed intracellular environment drives the evolution of a suite of distinct processes. These processes include meiotic chromosome segregation, ion homeostasis, and nucleotypic factors revolving around cell size, volume, and cell cycle control^7,10,11^. The broad array of relevant genes exhibiting signatures of selection in our data suggest that adaptations of each of these processes have a polygenic basis. Further, we observed the footprint of WGD-associated selection in genome-wide shifts in the frequencies of many alleles, replicated in three independent WGDs. This work, along with others^13,15–17^, shows that these processes exhibit genomic signatures of adaptive evolution consistent with observations of altered phenotype upon WGD consistently in independent adaptation events. A primary, well-established challenge occurs during meiotic chromosome segregation, and we here illustrated in *Cochlearia* other genomic and physiological changes. These physiological changes include increases in drought resilience and stomatal conductance, concordant with findings in other studies that concentrated on phenotype rather than genome-wide signal^10,82,83^.

Given this complex genomic architecture, we found that convergence in three species was minimal at the ortholog level. Even pairwise comparisons between species gave only a handful of common orthologs under selection (Table S9). However, at the level of functional processes, we observe evidence of convergent evolution. DNA management, as a high level process stood out, and in *Cochlearia* we saw a substantial shift relative to *A. arenosa:* in *Cochlearia* we observe enrichment in DNA repair and later meiotic processes, as opposed to the prophase I-oriented signal previously reported in *A. arenosa^3^.* It is not yet clear why particular solutions are favored in one species relative to another. A degree of stochasticity depending on available standing variation can be expected. But we also expect that an important role may also be played by difference in species histories, which may offer preadaptations that steer evolution in a particular direction. For example, our analysis of salinity tolerance in *Cochlearia* provided the surprising result the diploid cytotype was at least as tolerant to extreme levels of salt (600 mM, seawater concentrations) as the tetraploid, even though the diploid is found exclusively inland, with the tetraploids in seawater or directly on coasts. This cryptic salinity adaptation may be related to genetically connected polyploid coastal populations along the Atlantic coast from Portugal towards coastal systems in northern Scandinavia and the UK with glacial coastal refugia in the southern regions of Europe^44,84^. Furthermore diploid coastal populations from salt-affected habitat occur in Spain and a postglacial and boreal spread of the diploid towards the UK is possible^39, new^, with salinity tolerance developing along the way, altering the genomic substrate upon which selection acted in response to WGD-associated ionomic challenge^12,13^.

While we have not directly addressed the cause of the commonly observed adaptability of polyploids in this work, our results may suggest a hypothesis for one potential contributing factor to the occasional dramatic niche shifts observed following WGD. We observed a large quantity of DNA metabolism and repair genes under selection in all species, and especially *Cochlearia.* This may signal a temporarily increased susceptibility to DNA damage, due to suboptimal function of DNA repair genes during the process of adaptive evolution, resulting in a relative ‘mutator phenotype’ in young polyploids. Such a mutator phenotype has been plainly observed in aggressive polyploid human cancers, which not only exhibit SNP-level hypermutator phenotypes, but also dramatic structural variation in malignant aneuploid swarms that are associated with cancer progression^7^. It could be that a parallel to this exists following other WGD events, even in plants. Whether or not this hypothesis is supported by future discoveries, the cross-kingdom importance of WGD to fields from evolution, to ecology, to agriculture and medicine, underscores the importance of understanding the processes mediating adaptation to—and perhaps by—WGD.

## Methods

### Plant material

We first located 89 populations throughout Europe and collected population samplings of plants from each, aiming for at least 10 plants per population, with each sampled plant a minimum of 2 meters from any other. Of these, we selected 12 representative populations from British Isles and 12 populations from the rest of the European range for population resequencing (Table S3). An average of 4 individuals per population were sequenced with the exception of two hexaploid lineages that fall outside the central focus of this study, but were included for the demographic analysis, coastal *Cochlearia danica* (JON), which is invasive at inland stands, and coastal *Cochlearia anglica-like* FRE, for which only one individual was sequenced. A total of 116 individuals were initially sequenced, which was narrowed down by a cutoff of 4x, leaving 76 individuals from 23 populations in the final analysed dataset.

### Ploidy Determination

DNA content and ploidy were inferred for populations using Flow Cytometry (Dataset S1). Approximately 1 square cm of leaf material was diced alongside an internal reference using razor blades in 1 ml ice cold extraction buffer (either 45 mM MgCl_2_, 30 mM sodium citrate, 20mM MOPS, 1% Triton-100, pH 7 with NaOH for relative staining or 0.1 M citric acid, 0.5% Tween 20 for absolute measurements). The resultant slurry was then filtered through a 40-μm nylon mesh before the nuclei were stained with the addition of 1 ml staining buffer (either CyStain UV precise P [Sysmex, Fluorescence emission: 435nm to 500nm] for relative ploidy, or Otto 2 buffer [0.4 M Na_2_HPO_4_·12H_2_O, Propidium iodide 50 μgmL-1, RNase 50 μgmL-1], for absolute DNA content). After 1 minute of incubation at room temperature the sample was run for 5,000 particles on either a Partec PA II flow cytometer or a BD FACS Melody. Histograms were evaluated using FlowJo software version 10.6.1.

## Reference Genome Assembly and Alignment

We generated a synthetic long read-based *de novo* genome assembly using 10x Genomics Chromium linked read technology.

### CTAB DNA extraction method

A total of 0.4 g *Cochlearia pyrenaica* leaf material from one individual plant in the NEN population was ground using liquid nitrogen before the addition of 10 ml of CTAB DNA extraction buffer (100 mM Tris-HCl, 2% CTAB, 1.4 M NaCl, 20 mM EDTA, and 0.004 mg/ml Proteinase K). The mixture was incubated at 55°C for 1 hour then cooled on ice before the addition of 5 ml Chloroform. This was then centrifuged at 3000 rpm for 30 minutes and the upper phase taken, this was added to 1X volume of phenol:chloroform:isoamyl-alcohol and spun for 30 minutes at 3000 rpm. Again, the upper phase was taken and mixed with a 10% volume of 3M NaOAc and 2.5X volume of 100% ethanol at 4 °C. This was incubated on ice for 30 minutes before being centrifuged for 30 minutes at 3000 rpm and 4 °C. Three times the pellet was washed in 4ml 70% ethanol at 4 °C before being centrifuged again for 10 minutes at 3000 rpm and 4°C. The pellet was then air dried and resuspend in 300 ul nuclease-free water containing 0.0036 mg/ml RNase A. The DNA concentration was checked on a QuBit Flourometer 2.0 (Invitrogen) using the QuBit dsDNA HS Assay kit. Fragment sizes were assessed using a Q-card (OpGen Argus) and the Genomic DNA Tapestation assay (Agilent).

### 10X library construction

DNA material was diluted to 0.5 ng/μl with EB (Qiagen) and checked with a QuBit Flourometer 2.0 (Invitrogen) using the QuBit dsDNA HS Assay kit. The Chromium User Guide was followed as per the manufacturer’s instructions (10X Genomics, CG00043, Rev A). The final library was quantified using qPCR (KAPA Library Quant kit (Illumina), ABI Prism qPCR Mix, Kapa Biosystems). Sizing of the library fragments were checked using a Bioanalyzer (High Sensitivity DNA Reagents, Agilent). Samples were pooled based on the molarities calculated using the two QC measurements. The library was clustered at 8 pM with a 1 % spike of PhiX library (Illumina).

### Sequencing and assembly and assembly QC

The sample was sequenced on HiSeq2500 Rapid Run V2 mode (Illumina). The first run on 150 bp sequences gave 101.29 M reads. A second run was carried out on 250 bp sequences, bringing the total number of reads up to 269.58 M, total coverage to 123x and effective coverage to 51x. These were subsampled to 135 M reads and assembled on Supernova 2.0.0 giving raw coverage of 63x and effective coverage 35x. The molecule length was 16.6 kb. Two assemblies were kept, with a minimum contig size of 10 Kb or 3 Kb, with an assembly size of 174.47 Mb and 219.69 Mb respectively. The k-mer estimate for the genome size was 528.26 Mb and the flow cytometry estimate of the genome size was 656 Mb (0.67 pg +0.07) for the NEN diploid and 1,352 Mb (1.38 pg + 0.12) for the tetraploids, consistent with previous reports^85,86^. The final 3kb minimum contig length assembly had 13,302 contigs, an N50 of 39.7 kb. Assessment of gene space completeness gave 91.3% complete, single copy core eukaryotic ‘BUSCO genes’ (1315/1440 BUSCO groups; Table S1; BUSCO version 3.0.2).^87^ Uncollapsed haplotypes were detected in this assembly: to purge these we identified uncollapsed haplotypes (defined as ID > 99%, coverage > 99%; consisting ~28 mb of the genome) and one scaffold was randomly selected to use in alignments (consisting ~12 mb of the genome), while the rest were excluded.

### Gene Calling and Annotation

The genome was annotated with gene model predictions produced by AUGUSTUS (version 3)^88^ which had been trained on the *Arabidopsis thaliana* genome. A total of 44,023 putative genes were identified.

## Population Resequencing and Analysis

### Library preparation and sequencing

DNA was prepared using the commercially available DNeasy Plant Mini Kit from Qiagen. DNA libraries were made using TruSeq DNA PCR-free Library kit from Illumina as per the manufacturer’s instructions and were multiplexed based on concentrations measured with a QuBit Flourometer 2.0 (Invitrogen) using the QuBit dsDNA HS Assay kit. Sequencing was carried out on either NextSeq 550 (Illumina) in house (4 runs) or sent to Novogene for Illumina Hiseq X, PE150 sequencing (2 runs).

### Data preparation, alignment, and genotyping

Adapter sequences were removed using cutadapt (version 1.9.1)^89^ and quality trimmed with Sickle (version 1.2)^90^ to generate only high-quality reads (Phred score >=30) of 30bp or more. The reads were aligned to the reference using bwa (v. 0.7.12)^91^ and further processed with samtools (v. 1.3)^92^. Duplicate reads were removed and read group IDs added to the bam files using Picard (version 1.134)^93^. Indels were realigned with GATK (version 3.8)^94^. Samples were first genotyped individually with “HaplotypeCaller” and were then genotyped jointly using “GenotypeGVCFs” in GATK (version 3.8)^94^. The resulting VCF files were then filtered for biallelic sites and mapping quality (QD < 2.0, FS > 60.0, MQ < 40.0, MQRankSum < −12.5, ReadPosRankSum < −8.0, HaplotypeScore < 13.0). The VCF was then filtered by depth. To prevent a single individual from dominating the mean depths the three individuals with the most coverage (VEG_003, SKF_009 and VEG_004) were removed and a depth histogram was created for the remaining 73 individuals. Based on this distribution a depth cutoff of 2,469 was applied to the VCF containing the 73 individuals and this was then used as a mask for the final VCF containing all individuals.

### Demographic analysis

We inferred relationships between populations as genetic distances using principal component analysis (PCA) implemented in *adegenet*^95^. To further interrogate their relationships we then ran fastSTRUCTURE^96^. Since fastSTRUCTURE does not handle polyploid genomes we randomly subsampled two alleles from tetraploid and hexaploid populations using a custom script and used this dataset in fastStructure. We have previously demonstrated that results generated in this way are directly comparable to results generated with the full dataset in STRUCTURE^97^. We calculated Nei’s distances among all individuals in stamp and visualised these using SplitsTree^98^.

### Phylogenetic analysis

We constructed a maximum likelihood phylogeny using RAxML version 8.1.16^99^ under a GTR + G model of evolution and with an ascertainment bias correction (--asc-corr=lewis) in order to account for unsampled invariant sites in SNP datasets. Sites with more than 10% of missing data were excluded from a set of 79,252 fourfold-degenerate SNPs using the GATK tool “SelectVariants” (GATK version 3.8), and the python script ascbias.py (https://github.com/btmartin721/raxml_ascbias) was used to remove sites considered as invariable by RAxML. The maximum likelihood analysis was performed with 1000 rapid bootstrap replicates. Additionally, we used TreeMix version 1.13^45^ to generate a population maximum likelihood phylogeny allowing for migration events (admixture) between populations. The input file was generated using the script vcf2treemix.py (https://github.com/CoBiG2/RAD_Tools/blob/master/vcf2treemix.py), thereby excluding multiallelic sites from the set of fourfold-degenerate variants with a maximum of 10% missing data. We tested for up to 10 migration edges *(M)* and performed 10 initial replicate runs for every *M.* We then determined the optimal number of migration edges based on the Evanno method using the R package optM^100^. Hereafter, we performed 100 bootstrap replicates for the bestfitting *M* and finally, a consensus tree was inferred from the resulting 100 maximum likelihood trees using sumtrees.py version 4.10^101^.

### Orthogrouping and Reciprocal Best Blast Hits

We performed an orthogroup analysis using Orthofinder version 2.3.3^81^. to infer orthologous groups (OGs) from four species *(C. amara, A. lyrata, A. thaliana, C. pyrenaica).* A total of 21,618 OGs were found. Best reciprocal blast hits (RBHs) for *Cochlearia* and *A. thaliana* genes were found using BLAST version 2.9.0. *Cochlearia* genes were then assigned an *A. thaliana* gene ID for GO enrichment analysis in one of five ways. First if the genes’ OG contained only one *A. thaliana* gene ID, that gene ID was used. If the OG contained more than one *A. thaliana* gene ID then the RBH was taken. If there was no RBH then the OG gene with the lowest E-value in a BLAST versus the TAIR10 database was taken. If no OG contained the *Cochlearia* gene then the RBH was taken. Finally, if there was no OG or RBH then the gene with the lowest E-value in a BLAST versus the TAIR10 database was taken. BLASTs versus the TAIR10 database were performed during December 2019.

### GO Enrichment Analysis

To infer functions significantly associated with directional selection following WGD, we performed gene ontology enrichment of candidate genes in the R package TopGO v.2.32^102^, using *A. thaliana* orthologs of *Cochlearia* genes and an *A. thaliana* universe set. We tested for overrepresented Gene Ontology (GO) terms within the three domains Biological Process (BP), Cellular Component (CC) and Molecular Function (MF) using Fisher’s exact test with conservative ‘elim’ method, which tests for enrichment of terms from the bottom of the GO hierarchy to the top and discards any genes that are significantly enriched in a descendant GO term^103^. We used ‘biological process’ ontology with minimum node size 150 genes and FDR = 0.05. A significance cut-off of 0.05 and all processes represented by a single gene were removed (25 in total).

### Window-based scan for selective sweep signatures

We performed a window-based divergence scan for selection consisting of 1 kb windows that contained at least 20 SNPs. The data was filtered as described above and in addition was filtered for no more than 20% missing data and a depth of >= 8x. We calculated metrics: Rho, Nei’s Fst, Weir-Cochran’s Fst, FstH, Dxy, number of fixed differences and average groupwise allele frequency difference (AFD). To determine the best metric to use we performed a quantitative analysis of AFD plot quality for all 1% outliers of each metric. Each window was given a score of 0-4, with 0 being the lowest quality and 4 the highest. Scores were based on two qualities: peak height and peak specificity. For peak height one point was awarded if the window contained one SNP of AFD > 0.5 < 0.7, and two points were awarded for any SNP of AFD > 0.7. Likewise, for peak specificity two points were awarded for an AFD peak that was restricted to a single gene and one point was awarded for a peak that was restricted to 2-3 genes. The top 1% outliers from the metric FstH^47^ was selected as, compared to all other single 1% outlier lists and all permutations of overlapped 1% outlier lists, it maximized the number of ‘4’ and ‘3’ scores while minimizing the number of ‘1’ and ‘0’ scores. This is consistent with the good performance of Fst in our previous studies^12,13,17,52^.

### MAV analysis

A FineMAV^53^-like analysis was carried out on all biallelic, non-synonymous SNPs passing the same filters as the window-based selection scan. SNPs were assigned a Grantham score according to the amino acid change and this was scaled by the AFD between ploidies. The top 1% outliers of all these MAV-SNPs were then overlapped with the genes in our 1% Fst outlier windows to give a refined list of candidate genes that contain potentially functionally significant non-synonymous mutations at high AFD between cytotypes. The code outlining this can be found at https://github.com/paajanen/meiosis_protein_evolution/tree/master/FAAD.

## Supporting information

Dataset S1 Ploidy

Dataset S2 Cochlearia Fst

Dataset S3 Cochlearia MAV

Dataset S4 A_arenosa

Dataset S5 C_amara

## Data Availability

Sequence data that support the findings of this study have been deposited in the Sequence Read Archive (SRA; https://www.ncbi.nlm.nih.gov/sra) with the primary accession code PRJNAXXXXXX (available at http://www.ncbi.nlm.nih.gov/bioproject/XXXXX) and will be released following peer review and publication.

## Acknowledgements

The authors thank Kirsten Bomblies, Lara Hebberecht-Lopez, Filip Kolar, Mary Bray and Nigel Bray and for their assistance collecting plant material. This work was supported by the European Research Council (ERC) under the European Union’s Horizon 2020 research and innovation programme [grant number ERC-StG 679056 HOTSPOT], via a grant to LY.

## Author Contributions

LY and SMB conceived the study. SMB, EW, MB, SB, SF, PP, MK and LY performed analyses. SMB, SB, MZ, and SF performed laboratory experiments. LY, SMB, PM, and JK performed field collections. LY and SMB wrote the manuscript with primary input from all authors. All authors edited and approved the final manuscript.

## Materials & Correspondence

Correspondence and material requests should be addressed to Levi Yant at

