## Supplementary information for "Convergence and novelty in adaptation to whole genome duplication in three independent polyploids"

#### Supplementary Text

##### Supplementary Text 1: Salinity tolerance experiments.

Seeds were sowed in petri plates filled with wet sand for a week, then transferred to pot trays filled with John Innes mix (No.1). Each tray contained 60 plants. A total of 1,036 plants were planted. The experiment was conducted in complete randomized design with as many replications as possible per maternal seed line (average 13.05) and plants were grown in a glasshouse at ambient UK summer temperature (~25C day, ~10C night). After 2 weeks from germination, trays were bottom-watered twice weekly with watering solution containing either 10 mM MES pH 7 and NaCl for the treatment or just 10 mM MES for the control. Salt concentrations began at 50 mM NaCl for two weeks, then were increased to 300 mM for two weeks, and were finally increased to 600 mM for 4 weeks. During the final week, leaves were sprayed with 600 mM NaCl watering solution twice. After 8 total weeks of treatment, a modified Standard Evaluation System (SES) Score was used to evaluate symptoms of salt stress[1]. Three individuals independently took measurements for each plant and the final score was given as the average of the three. At the same time point, 2 leaves of each plant were harvested, dried at 60 °C for 2 days and stored for ICP analysis. The diploids were significantly more salt tolerant than the tetraploids (W = 29568, p-value = 2.178e-05, Wilcoxon rank sum test with continuity correction performed in R).

##### Supplementary Text 2: Stomatal conductance assessment, net photosynthesis and drought tolerance experiments.

To test for differences in physiological responses to drought stress we cultivated twenty different populations of *Cochlearia* side by side in a controlled environment. Seeds were germinated in wet sand in a growth camber with temperature and humidity control before being transplanted into PET-based potting soil and placed in a growth room (16h light from 6:00 to 20:00, temperature day: 22°C and night: 18°C). The watering regime for control and drought stressed plants was as follows: For the first 2 weeks after transplanting, they were watered 1000 ml $H_2O$ per tray every 2$^{nd}$ day. From the 3$^{rd}$ week onwards a drought stress was imposed. Control plants were still watered with 1000 ml $H_2O$ per tray every 2$^{nd}$ day. Drought stress was generated by lowering watering/soil moisture week by week as follows: week 3 - 1000 ml $H_2O$ per tray every 3$^{rd}$ day, week 4 – 1000 ml $H_2O$ per tray every 4$^{th}$ day, week 5 – 1000 ml $H_2O$ per tray every 5$^{th}$ day, week 6 – 1000 ml $H_2O$ per tray every 6$^{th}$ day. Plants were assessed at the end of week 6. Wet and dry weight of the shoot as well as photosynthesis parameters were captured. Photosynthetic parameters were assessed using the LI-6400XT

Portable Photosynthesis System (LI-COR, Lincoln, NE, United States). All measurements were taken between 9:30 and 11:30 using light adapted plants. Photosynthetic parameters were measured on the largest leaf from both control and drought treatment groups and all measurements were conducted at the same time of the day (9:30am-11:30am). The leaf was placed into a custom-made single leaf chamber (chamber: polyoxymethylene, lid: poly (methyl methacrylate)), openings sealed with sponge rubber, which was connected to the LI-COR. Photosynthetic parameters were recorded after the reads were stable. We analysed 9 diploid populations and 11 tetraploid populations. We observed an increased net photosynthetic rate in tetraploids in both control and drought conditions (Figure S1 and Table S6). We note large population-specific variation in both cytotypes.

Figure S1 and Table S7 show stomatal conductance (SC) of diploid and tetraploid populations under drought stress relative to well-watered conditions (relative SC). Averages, standard deviation and confidence interval for relative stomatal conductance and net photosynthesis can be found in Tables S6 and S7. Tetraploids show higher SC under drought stress. As the linear model in Table S8 shows, ploidy significantly affects the phenotype.

Data were analysed using R Version 3.6.0 (2019-04-26) with RStudio Version 1.2.1335. Data were plotted using the ggplot2 package version 3.2.1. The lsmeans package version 2.30-0 was used to fit a linear model to the data while the package bestNormalize version 1.4.3 was used for data normalization and estimation of normality. The package sjPlot version 2.8.2 was used to generate a summary table for the linear model fitted to the data. The package flextable version 0.5.9 was used to generate summary tables containing averages, sd and ci intervals.

1. Gregoria, G. B., Senadhira, D. & Mendoza, R. D. *Screening rice for salinity tolerance*. https://econpapers.repec.org/RePEc:ags:irridp:287589 (1997).

#### Supplemental Figure

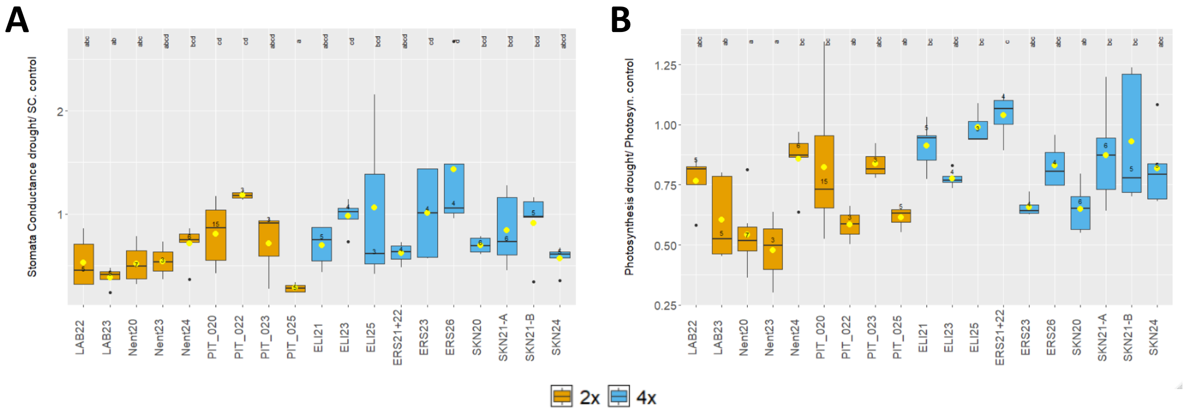


**Figure S1**. Ploidy effect on stomatal conductance and drought tolerance. Box plots show median and variation of **A)** stomatal conductance and **B)** net photosynthetic rate of drought stressed plants in comparison to well-watered plants. One-Way ANOVA, Post-hoc Tukey, n: as indicated above boxes, yellow dot: mean.

#### Supplemental Tables

**Table S1.** Assembly metrics of *Cochlearia* linked-read assemblies.

| Contig Minimum Size Cutoff | #contigs | L50 | N80 | N50 | Mean | Max | Sum |
|---|---|---|---|---|---|---|---|
| **10kb** | 4742 | 946 | 22,645 | 53,194 | 75,345 | 845,631 | 1.716E+08 |
| **3Kb** | 13302 | 1434 | 9,568 | 39,689 | 60,908 | 845,631 | 2.167E+08 |

**Table S2.** BUSCO metrics giving estimate of gene space completeness.

| Minimum contig size cutoff: | 10 Kb | 3 Kb |
|---|---|---|
| Complete BUSCOs | 1254 (87.1%) | 1315 (91.3%) |
| Complete and Single-copy BUSCOS | 1135 (78.8%) | 1178 (81.8%) |
| Complete amd duplicated BUSCOs | 119 (8.3%) | 137 (9.5%) |
| Fragmented BUSCOs | 33 (2.3%) | 45 (3.1%) |
| Missing BUSCOs | 153 (10.6%) | 80 (5.6%) |
| Total BUSCO groups searched | 1440 (100%) | 1440 (100%) |

**Table S3.** Sequenced individuals, sequencing depth and locality information.

| Population | Number of plants | Names | Mean Depth (coverage) | Ploidy | Putative Species | Latitude | Longitude | Country | Locality |
|---|---|---|---|---|---|---|---|---|---|
| **ALO** | 4 | ALO_006 | 14.50 | 4 | *C. officinalis* | 56.1028 | -3.80791 | UK | Riverside (estuary) |
| | | ALO_007 | 16.08 | | | | | | |
| | | ALO_013 | 24.74 | | | | | | |
| | | ALO_017 | 40.91 | | | | | | |
| **BAQ** | 0 | *BAQ_001* | *1.11* | 2 | *C. pyrenaica* | 42.70325 | 0.94767 | Spain | Spring |
| | | *BAQ_002* | *1.28* | | | | | | |
| | | *BAQ_003* | *1.26* | | | | | | |
| | | *BAQ_004* | *1.3* | | | | | | |
| **BEA** | 3 | *BEA_001* | *1.04* | 4 | *C. officinalis* | 69.39424 | 19.45429 | Norway | Sea |
| | | BEA_002 | 24.70 | | | | | | |
| | | BEA_004 | 6.59 | | | | | | |
| | | *BEA_007* | *3.99* | | | | | | |
| | | *BEA_009* | *1.02* | | | | | | |
| | | BEA_010 | 9.83 | | | | | | |
| **BRI** | 4 | *BRI_001* | *1.03* | 4 | *C. officinalis* | 69.69288 | 18.88216 | Norway | Sea |
| | | BRI_002 | 14.56 | | | | | | |
| | | *BRI_004* | *1.02* | | | | | | |
| | | BRI_005 | 33.66 | | | | | | |
| | | BRI_006 | 7.23 | | | | | | |
| | | BRI_009 | 8.42 | | | | | | |
| **ELI** | 4 | ELI_001 | 19.15 | 4 | *C. officinalis* | 56.18478 | -2.81303 | UK | Sea |
| | | ELI_002 | 17.59 | | | | | | |
| | | ELI_003 | 21.93 | | | | | | |
| | | ELI_004 | 19.56 | | | | | | |
| **ESH** | 0 | *ESH_004* | *1.29* | 4 | *C. officinalis* | 60.48972 | -1.62806 | UK | Coastal cliffs (bird nests) |
| | | *ESH_005* | *1.12* | | | | | | |
| | | *ESH_006* | *1.18* | | | | | | |
| | | *ESH_100* | *1.14* | | | | | | |
| **FRE** | 1 | FRE_013 | 22.63 | 6 | *C. anglica-like* | 51.07818 | -4.12301 | UK | Salt marsh |
| **JON** | 1 | JON_001 | 14.47 | 6 | *C. danica* | 52.62162 | 1.22366 | UK | Road grit |
| **KVA** | 4 | KVA_002 | 19.41 | 4 | *C. officinalis* | 69.6888 | 18.83254 | Norway | Spring |
| | | KVA_003 | 20.34 | | | | | | |
| | | *KVA_006* | *1.03* | | | | | | |
| | | *KVA_008* | *1.02* | | | | | | |
| | | KVA_009 | 6.70 | | | | | | |
| | | KVA_010 | 25.99 | | | | | | |
| | | *KVA_101* | *1.01* | | | | | | |
| **LAB** | 4 | LAB_004 | 26.96 | 2 | *C. pyrenaica* | 54.67205 | -2.231 | UK | Riverside (inland) |
| | | LAB_300 | 14.49 | | | | | | |
| | | LAB_400 | 17.33 | | | | | | |
| | | LAB_500 | 27.30 | | | | | | |

| | | | | | | | | | |
|---|---|---|---|---|---|---|---|---|---|
| **LNL** | 4 | LNL_001 | 19.05 | 4 | *C. micacea* | 56.53804 | -4.29126 | UK | Crags along the banks of a dam |
| | | LNL_002 | 16.37 | | | | | | |
| | | LNL_003 | 19.72 | | | | | | |
| | | LNL_008 | 18.15 | | | | | | |
| **MEL** | 2 | MEL_001 | 14.09 | 4 | *C. officinalis* | 69.26708 | 19.92057 | Norway | Sea |
| | | MEL_002 | 7.78 | | | | | | |
| | | *MEL_003* | *1.02* | | | | | | |
| | | *MEL_004* | *1.02* | | | | | | |
| **NEN** | 4 | NEN_001 | 20.20 | 2 | *C. pyrenaica* | 54.81497 | -2.43521 | UK | Riverside (inland) |
| | | NEN_003 | 20.19 | | | | | | |
| | | NEN_200 | 17.80 | | | | | | |
| | | NEN_300 | 18.67 | | | | | | |
| **ROT** | 4 | ROT_004 | 16.59 | 4 | *C. officinalis* | 54.49043 | -3.60484 | UK | Sea |
| | | ROT_006 | 19.10 | | | | | | |
| | | ROT_007 | 36.28 | | | | | | |
| | | ROT_013 | 30.38 | | | | | | |
| **RUZ** | 1 | RUZ_001 | 10.59 | 2 | *C. pyrenaica* | 49.07523 | 19.3066 | Slovakia | Travertine spring |
| | | *RUZ_004* | *1.04* | | | | | | |
| **SAL** | 3 | *SAL_001* | *1.02* | 2 | *C. pyrenaica* | 48.12515 | 12.75423 | Germany | Spring rivulet |
| | | SAL_003 | 21.37 | | | | | | |
| | | SAL_004 | 19.22 | | | | | | |
| | | *SAL_005* | *14.80* | | | | | | |
| | | SAL_006 | 9.83 | | | | | | |
| | | *SAL_007* | *1.02* | | | | | | |
| | | *SAL_008* | *1.02* | | | | | | |
| **SKF** | 4 | SKF_002 | 7.07 | 6 | *C. anglica-like* | 53.64328 | 0.06969 | UK | Sea |
| | | SKF_003 | 19.60 | | | | | | |
| | | SKF_005 | 32.47 | | | | | | |
| | | SKF_009 | 80.70 | | | | | | |
| | | *SKF_013* | *1.02* | | | | | | |
| | | *SKF_015* | *1.06* | | | | | | |
| **SKI** | 3 | SKI_004 | 5.02 | 4 | *C. officinalis* | 69.37866 | 20.23541 | UK | Riverside (estuary) |
| | | SKI_005 | 25.51 | | | | | | |
| | | *SKI_006* | *1.03* | | | | | | |
| | | *SKI_007* | *1.11* | | | | | | |
| | | SKI_008 | 4.25 | | | | | | |
| | | *SKI_009* | *1.03* | | | | | | |
| | | *SKI_010* | *1.02* | | | | | | |
| **SKN** | 4 | SKN_001 | 15.96 | 4 | *C. officinalis* | 54.57107 | -0.8962 | UK | Sea |
| | | SKN_002 | 16.51 | | | | | | |
| | | SKN_005 | 31.30 | | | | | | |
| | | SKN_008 | 27.77 | | | | | | |
| **SPU** | 4 | *SPU_004* | *1.03* | 2 | *C. pyrenaica* | 53.60964 | 0.1438 | UK | Sea |
| | | SPU_006 | 11.72 | | | | | | |
| | | SPU_008 | 16.77 | | | | | | |

| | | | | | | | | | |
|---|---|---|---|---|---|---|---|---|---|
| | | SPU_009 | 10.88 | | | | | | |
| | | SPU_010 | 8.56 | | | | | | |
| **TET** | 4 | TET_002 | 15.31 | 4 | *C. officinalis* | 53.52525 | 0.01828 | UK | Sea |
| | | TET_004 | 22.65 | | | | | | |
| | | TET_006 | 14.53 | | | | | | |
| | | TET_008 | 13.17 | | | | | | |
| **TRO** | 4 | TRO_001 | 15.91 | 4 | *C. officinalis* | 69.62492 | 19.04544 | Norway | Spring rivulet |
| | | *TRO_002* | *1.08* | | | | | | |
| | | TRO_003 | 11.73 | | | | | | |
| | | TRO_005 | 36.02 | | | | | | |
| | | *TRO_006* | *1.04* | | | | | | |
| | | *TRO_007* | *1.01* | | | | | | |
| | | *TRO_008* | *1.02* | | | | | | |
| | | TRO_009 | 23.30 | | | | | | |
| **VAG** | 4 | *VAG_001* | *1.02* | 4 | *C. officinalis* | 69.77513 | 19.29306 | Norway | Sea |
| | | VAG_003 | 10.97 | | | | | | |
| | | VAG_004 | 25.55 | | | | | | |
| | | VAG_007 | 35.57 | | | | | | |
| | | *VAG_008* | *1.01* | | | | | | |
| | | *VAG_010* | *1.05* | | | | | | |
| | | VAG_102 | 6.82 | | | | | | |
| | | *VAG_105* | *1.01* | | | | | | |
| **VEG** | 2 | *VEG_001* | *1.46* | 4 | *C aestuaria* | 43.46531 | -7.05268 | Spain | Riverside (urban) |
| | | *VEG_002* | *1.2* | | | | | | |
| | | VEG_003 | 117.01 | | | | | | |
| | | VEG_004 | 62.03 | | | | | | |
| **WOL** | 4 | WOL_002 | 24.32 | 2 | *C. pyrenaica* | 47.8316 | 9.76181 | Germany | Spring |
| | | WOL_006 | 18.24 | | | | | | |
| | | WOL_009 | 20.40 | | | | | | |
| | | WOL_010 | 18.96 | | | | | | |

**Note:** The second column indicates the number of individuals from each population in the analysis, a total of 76. Individuals whose name and depth is in grey were excluded due to sequencing depth or suspected contamination (SAL_005 only).

**Table S4.** GO Enrichment among selective sweep candidates in *Cochlearia*.

| GO.ID | Genes | Term | Annotated | Significant | Expected | Fisher |
|---|---|---|---|---|---|---|
| **GO:0051301** | **AT1G15660, AT1G80080, AT2G14120, AT3G10220, AT3G12280, AT3G22780, AT3G55340, AT4G21270, AT4G32410, AT5G15540, AT5G22220, AT5G44030, AT5G60410** | **cell division** | **435** | **13** | **4.32** | **0.00044** |
| **GO:0006997** | AT1G19520, AT2G05120, AT3G10650, AT5G51200, AT5G56210 | nucleus organization | 71 | 5 | 0.71 | 0.00071 |
| **GO:0000819** | AT1G15660, AT3G16730, AT3G10440, AT4G05360, AT5G10010, AT5G15540, AT5G37630 | sister chromatid segregation | 93 | 7 | 0.92 | 0.00131 |
| **GO:0070192** | AT3G16730, AT3G10440, AT3G12280, AT5G15540 | chromosome organization involved in meiotic cell cycle | 50 | 4 | 0.5 | 0.00153 |
| **GO:0045132** | AT1G15660, AT3G10440, AT3G12280, AT5G15540 | meiotic chromosome segregation | 61 | 4 | 0.61 | 0.00318 |
| **GO:0034508** | AT1G15660, AT5G15540 | centromere complex assembly | 11 | 2 | 0.11 | 0.00509 |
| **GO:0045144** | AT3G10440, AT5G15540 | meiotic sister chromatid segregation | 16 | 2 | 0.16 | 0.01075 |
| **GO:0051177** | AT3G10440, AT5G15540 | meiotic sister chromatid cohesion | 16 | 2 | 0.16 | 0.01075 |
| **GO:0007062** | AT3G10440, AT4G05360, AT5G15540 | sister chromatid cohesion | 49 | 3 | 0.49 | 0.01273 |
| **GO:0007049** | **AT1G15660, AT2G14120, AT2G32710, AT3G07100, AT3G10440, AT3G12280, AT3G16730, AT3G22780, AT3G50360, AT3G55340, AT4G05360, AT4G21270, AT4G32410, AT4G35520, AT5G10010, AT5G15540, AT5G23860, AT5G22220, AT5G37630, AT5G44030, AT5G45400** | **cell cycle** | **753** | **21** | **7.48** | **0.01500** |
| **GO:0006310** | **AT3G05850, AT4G23160, AT4G35520, AT5G15540, AT5G45400, AT5G57160** | **DNA recombination** | **201** | **6** | **2** | **0.01543** |
| **GO:0022402** | **AT1G15660, AT2G32710, AT3G07100, AT3G10440, AT3G12280, AT3G16730, AT3G22780, AT3G50360, AT3G55340, AT4G05360, AT4G21270, AT4G32410, AT4G35520, AT5G10010, AT5G15540, AT5G37630, AT5G44030** | **cell cycle process** | **508** | **17** | **5.04** | **0.01640** |
| **GO:0006259** | **AT1G06670, AT1G48720, AT1G67020, AT2G02090, AT3G05850, AT3G07100, AT3G12280, AT3G16980, AT3G16730, AT3G31430, AT4G00980, AT4G04650, AT4G05360, AT4G23160, AT4G29090, AT4G35520, AT5G03450, AT5G15540, AT5G42905, AT5G45400, AT5G57160, ATMG00300, ATMG00860, ATMG00750, ATMG00810** | **DNA metabolic process** | **742** | **25** | **7.37** | **0.01764** |
| **GO:0006928** | AT1G21730, AT2G33240, AT3G10180, AT4G21270 | movement of cell or subcellular component | 102 | 4 | 1.01 | 0.01890 |
| **GO:0007135** | AT3G10440, AT5G15540 | meiosis II | 23 | 2 | 0.23 | 0.02166 |
| **GO:0061983** | AT3G10440, AT5G15540 | meiosis II cell cycle process | 23 | 2 | 0.23 | 0.02166 |

| | | | | | | |
|---|---|---|---|---|---|---|
| **GO:0090304** | **AT1G09220, AT5G57160, AT2G22670, AT3G06380, AT4G23160, AT5G04670, AT3G16980, AT3G16730, AT3G31430, AT3G05850, AT3G07810, ATMG00860, AT4G13650, ATMG00300, AT4G05360, AT4G35520, ATMG00750, AT1G49900, AT5G37190, AT5G42905, AT4G16845, AT1G67020, AT1G06670, AT1G06740, AT5G22650, AT5G15690, AT2G46680, AT1G48720, ATMG00810, AT4G13040, AT4G00980, AT4G10070, AT4G08540, AT5G24120, AT1G50200, AT4G24650, AT4G29090, AT5G64270, AT2G05120, AT1G64490, AT3G22780, AT2G25900, AT5G16780, AT5G16715, AT4G14850, AT2G19910, AT5G11430, AT3G62310, AT5G64420, AT1G55870, AT1G55325, AT1G72090, AT3G61050, AT5G22220, AT3G12270, AT3G12280, AT1G69600, AT2G02980, AT4G04650, AT3G25660, AT2G46510, AT5G58850, AT5G45400, AT5G03450, AT5G15540, AT3G07100, AT2G02090, AT2G01370, AT3G17609** | **nucleic acid metabolic process** | **4577** | **69** | **45.45** | **0.02961** |
| **GO:0000280** | **AT1G15660, AT3G10440, AT3G12280, AT3G16730, AT3G55340, AT4G21270, AT4G35520, AT5G10010, AT5G15540, AT5G37630** | **nuclear division** | **251** | **10** | **2.49** | **0.03186** |
| **GO:0008356** | AT1G80080, AT3G12280 | asymmetric cell division | 30 | 2 | 0.3 | 0.03559 |
| **GO:0007127** | AT1G15660, AT3G12280, AT4G35520 | meiosis I | 75 | 3 | 0.74 | 0.03873 |
| **GO:0051321** | **AT1G15660, AT3G10440, AT3G12280, AT3G16730, AT4G21270, AT4G35520, AT5G15540, AT5G22220, AT5G45400** | **meiotic cell cycle** | **197** | **9** | **1.96** | **0.03950** |
| **GO:0045143** | AT1G15660, AT3G12280 | homologous chromosome segregation | 33 | 2 | 0.33 | 0.04238 |
| **GO:0061982** | AT1G15660, AT3G12280, AT4G35520 | meiosis I cell cycle process | 81 | 3 | 0.8 | 0.04689 |
| **GO:0048609** | **AT2G39940, AT3G07100, AT4G21270, AT4G39010, AT5G22650** | **multicellular organismal reproductive pr...** | **197** | **5** | **1.96** | **0.04718** |
| **GO:0033045** | AT5G10010, AT5G15540 | regulation of sister chromatid segregati... | 36 | 2 | 0.36 | 0.04960 |
| **GO:0015074** | AT1G67020, AT1G48720, AT4G23160, AT4G00980, ATMG00860, ATMG00300, ATMG00750, ATMG00810 | DNA integration | 26 | 8 | 0.26 | 1.1E-10 |
| **GO:0006278** | AT1G48720, AT3G31430, AT4G04650, AT4G23160, AT4G29090, AT5G42905, AT5G45400, ATMG00860, ATMG00300 | RNA-dependent DNA biosynthetic process | 87 | 9 | 0.86 | 2.1E-07 |
| **GO:0106074** | AT3G05850, AT1G50200, AT5G16715 | aminoacyl-tRNA metabolism involved in translational fidelity | 16 | 3 | 0.16 | 0.00049 |
| **GO:0010218** | AT2G39940, AT3G17609, AT4G10340, AT5G24120 | response to far red light | 55 | 4 | 0.55 | 0.00217 |
| **GO:0006561** | AT5G14800, AT2G39800 | proline biosynthetic process | 8 | 2 | 0.08 | 0.00264 |
| **GO:0010374** | AT1G80080, AT2G40180, AT3G22780, AT3G12280 | stomatal complex development | 63 | 4 | 0.63 | 0.00357 |
| **GO:0009651** | **AT1G09210, AT1G11910, AT1G55870, AT2G01980, AT2G39800, AT3G61050, AT4G10310, AT4G32410, AT5G08620, AT5G14800, AT5G19320, AT5G23860, AT5G24120, AT5G35630** | **response to salt stress** | **620** | **14** | **6.16** | **0.00379** |

| | | | | | | |
|---|---|---|---|---|---|---|
| **GO:0010090** | AT1G80080, AT3G12280, AT5G22220, AT5G65530 | trichome morphogenesis | 67 | 4 | 0.67 | 0.00445 |
| **GO:0007076** | AT5G37630, AT5G15540 | mitotic chromosome condensation | 11 | 2 | 0.11 | 0.00509 |
| **GO:0048285** | **AT1G15660, AT1G58200, AT2G14120, AT3G10440, AT3G12280, AT3G16730, AT3G55340, AT4G21270, AT4G35520, AT5G10010, AT5G15540, AT5G37630** | **organelle fission** | **295** | **12** | **2.93** | **0.00656** |
| **GO:0006349** | AT4G16845, AT3G12280 | regulation of gene expression by genetic imprinting | 13 | 2 | 0.13 | 0.00713 |
| **GO:0035725** | AT2G01980, AT4G10310 | sodium ion transmembrane transport | 13 | 2 | 0.13 | 0.00713 |
| **GO:0098659** | AT1G18910, AT2G01980 | inorganic cation import across plasma membrane | 13 | 2 | 0.13 | 0.00713 |
| **GO:1903047** | **AT1G15660, AT3G16730, AT3G12280, AT3G22780, AT3G55340, AT4G32410, AT5G10010, AT5G15540, AT5G37630, AT5G44030** | **mitotic cell cycle process** | **299** | **10** | **2.97** | **0.00797** |
| **GO:0009553** | AT1G19520, AT3G12280, AT5G24120, AT5G55820, AT5G60410 | embryo sac development | 146 | 5 | 1.45 | 0.01543 |
| **GO:0000281** | AT3G22780, AT3G55340, AT4G32410, AT5G44030 | mitotic cytokinesis | 96 | 4 | 0.95 | 0.01547 |
| **GO:0006281** | **AT2G02090, AT3G16980, AT3G16730, AT4G05360, AT4G35520, AT5G03450, AT5G15540, AT5G45400, AT5G57160** | **DNA repair** | **387** | **9** | **3.84** | **0.01568** |
| **GO:0010337** | AT4G13040, AT5G60410 | regulation of salicylic acid metabolic process | 20 | 2 | 0.2 | 0.01659 |
| **GO:0010440** | AT2G40180, AT3G22780 | stomatal lineage progression | 20 | 2 | 0.2 | 0.01659 |
| **GO:0010286** | AT4G17250, AT5G10010, AT5G60410 | heat acclimation | 56 | 3 | 0.56 | 0.01821 |
| **GO:0061640** | AT3G22780, AT3G55340, AT4G32410, AT5G44030 | cytoskeleton-dependent cytokinesis | 101 | 4 | 1 | 0.01830 |
| **GO:0002831** | AT1G80080, AT3G55150, AT4G13040, AT5G60410 | regulation of response to biotic stimulus | 103 | 4 | 1.02 | 0.01952 |
| **GO:0000070** | AT1G15660, AT3G16730, AT5G10010, AT5G37630, AT5G15540 | mitotic sister chromatid segregation | 69 | 5 | 0.69 | 0.01956 |
| **GO:0006813** | AT2G01980, AT4G04850, AT4G10310, AT4G19960 | potassium ion transport | 106 | 4 | 1.05 | 0.02144 |
| **GO:0006289** | AT3G16980, AT5G57160, AT5G45400 | nucleotide-excision repair | 60 | 3 | 0.6 | 0.02185 |
| **GO:0030003** | **AT1G18910, AT2G01980, AT2G18960, AT5G36290, AT5G52790** | **cellular cation homeostasis** | **160** | **5** | **1.59** | **0.02199** |
| **GO:0000910** | AT3G22780, AT3G55340, AT4G32410, AT5G44030 | cytokinesis | 108 | 4 | 1.07 | 0.02278 |
| **GO:0032101** | AT1G80080, AT3G55150, AT4G13040, AT5G60410 | regulation of response to external stimulus | 108 | 4 | 1.07 | 0.02278 |
| **GO:0009084** | AT2G39800, AT5G14800, AT4G17830, AT5G35630 | glutamine family amino acid biosynthetic process | 33 | 4 | 0.33 | 0.02496 |

| | | | | | | |
|---|---|---|---|---|---|---|
| **GO:0006974** | **AT2G02090, AT3G16980, AT3G16730, AT4G05360, AT4G35520, AT5G03450, AT5G15540, AT5G45400, AT5G57160** | **cellular response to DNA damage stimulus** | **420** | **9** | **4.17** | **0.02504** |
| **GO:0006020** | AT1G14520, AT5G16760 | inositol metabolic process | 25 | 2 | 0.25 | 0.02535 |
| **GO:1900426** | AT3G55150, AT4G13040 | positive regulation of defense response to bacterium | 25 | 2 | 0.25 | 0.02535 |
| **GO:0071804** | AT2G01980, AT4G10310, AT4G19960 | cellular potassium ion transport | 64 | 3 | 0.64 | 0.02585 |
| **GO:0071805** | AT2G01980, AT4G10310, AT4G19960 | potassium ion transmembrane transport | 64 | 3 | 0.64 | 0.02585 |
| **GO:0043044** | AT2G02090, AT3G12810 | ATP-dependent chromatin remodeling | 26 | 2 | 0.26 | 0.02729 |
| **GO:0050777** | AT5G60410, AT5G35580 | negative regulation of immune response | 26 | 2 | 0.26 | 0.02729 |
| **GO:0006873** | **AT1G18910, AT2G01980, AT2G18960, AT5G36290, AT5G52790** | **cellular ion homeostasis** | **172** | **5** | **1.71** | **0.02886** |
| **GO:0043038** | AT1G50200, AT3G25660, AT5G16715 | amino acid activation | 67 | 3 | 0.67 | 0.02909 |
| **GO:0043039** | AT1G50200, AT3G25660, AT5G16715 | tRNA aminoacylation | 67 | 3 | 0.67 | 0.02909 |
| **GO:0009833** | AT4G32410, AT5G44030 | plant-type primary cell wall biogenesis | 27 | 2 | 0.27 | 0.02928 |
| **GO:0009900** | AT2G39940, AT4G39010 | dehiscence | 27 | 2 | 0.27 | 0.02928 |
| **GO:0010207** | AT4G10340, AT5G24120 | photosystem II assembly | 27 | 2 | 0.27 | 0.02928 |
| **GO:0002683** | AT5G60410, AT5G35580 | negative regulation of immune system process | 28 | 2 | 0.28 | 0.03133 |

| | | | | | | |
|---|---|---|---|---|---|---|
| **GO:0043170** | **AT1G09220, AT1G11480, AT1G06670, AT1G06740, AT1G18910, AT1G11910, AT1G28440, AT2G22670, AT3G06380, AT4G23160, AT3G16980, AT3G16730, AT3G31430, AT3G05850, AT3G22750, AT3G07810, AT4G13650, AT4G05360, AT1G49900, AT3G57760, AT4G16845, AT1G67020, AT2G06105, AT3G62010, AT3G22180, AT2G46680, AT1G48720, AT4G10340, AT4G11110, AT4G13040, AT3G12810, AT3G47890, AT4G00980, AT4G20790, AT2G40180, AT3G61740, AT4G10070, AT4G08540, AT1G50200, AT4G24650, AT2G26140, AT2G05120, AT2G32710, AT1G64490, AT3G22780, AT2G25900, AT4G14850, AT2G43050, AT2G19910, AT3G62310, AT3G17420, AT1G55870, AT1G55325, AT1G72090, AT1G60995, AT3G61050, AT2G19210, AT3G12270, AT3G12280, AT1G69830, AT1G69600, AT2G02980, AT4G04650, AT3G10650, AT3G25660, AT3G05790, AT2G46510, AT4G14320, AT2G39940, AT3G07100, AT2G02090, AT2G01370, AT3G17609, AT5G57160, AT5G04670, AT5G02880, ATMG00860, AT4G29990, ATMG00300, AT5G25510, AT5G35980, AT4G35520, ATMG00750, AT5G37190, AT5G42905, AT5G45570, AT5G22650, AT5G15690, AT5G07270, ATMG00810, AT5G60410, AT5G24120, AT4G29090, AT5G64270, AT5G35580, AT4G31160, AT5G65530, AT4G32410, AT5G16780, AT5G16715, AT5G16590, AT5G11430, AT5G64420, AT5G42620, AT5G22220, AT5G14260, AT5G51770, AT5G19180, AT5G44030, AT5G58850, AT5G45400, AT5G10010, AT5G03450, AT4G39010, AT5G15540** | **macromolecule metabolic process** | **9179** | **115** | **91.15** | **0.03246** |
| **GO:0007018** | AT1G21730, AT3G10180, AT4G21270 | microtubule-based movement | 70 | 3 | 0.7 | 0.03253 |
| **GO:2000142** | AT1G64490, AT5G24120 | regulation of DNA-templated transcription, initiation | 29 | 2 | 0.29 | 0.03344 |
| **GO:0006913** | AT2G05120, AT3G10650, AT5G19320, AT5G51200 | nucleocytoplasmic transport | 122 | 4 | 1.21 | 0.03358 |
| **GO:0051169** | AT2G05120, AT3G10650, AT5G19320, AT5G51200 | nuclear transport | 122 | 4 | 1.21 | 0.03358 |
| **GO:0010114** | AT3G17609, AT4G10340, AT5G24120 | response to red light | 72 | 3 | 0.72 | 0.03494 |
| **GO:0010099** | AT3G17609, AT5G37190 | regulation of photomorphogenesis | 30 | 2 | 0.3 | 0.03559 |
| **GO:0010103** | AT1G80080, AT3G12280 | stomatal complex morphogenesis | 30 | 2 | 0.3 | 0.03559 |
| **GO:0010629** | **AT2G02090, AT2G19910, AT2G22670, AT2G25900, AT2G46510, AT3G12810, AT3G12280, AT3G61050, AT5G10010, AT5G22650, AT5G22220** | **negative regulation of gene expression** | **597** | **11** | **5.93** | **0.03691** |
| **GO:0071215** | **AT1G80080, AT2G40180, AT2G46510, AT3G06380, AT3G50360, AT5G35980, AT5G60410** | **cellular response to abscisic acid stimu...** | **313** | **7** | **3.11** | **0.03738** |
| **GO:0097306** | **AT1G80080, AT2G40180, AT2G46510, AT3G06380, AT3G50360, AT5G35980, AT5G60410** | **cellular response to alcohol** | **313** | **7** | **3.11** | **0.03738** |

| | | | | | | |
|---|---|---|---|---|---|---|
| **GO:0042364** | AT1G14520, AT2G46580, AT3G16990 | water-soluble vitamin biosynthetic proce... | 74 | 3 | 0.73 | 0.03744 |
| **GO:0051028** | AT2G05120, AT3G10650, AT5G51200 | mRNA transport | 74 | 3 | 0.73 | 0.03744 |
| **GO:0033047** | AT5G10010, AT5G15540 | regulation of mitotic sister chromatid s... | 31 | 2 | 0.31 | 0.03780 |
| **GO:0032392** | AT1G06670, AT3G12810, AT5G45400 | DNA geometric change | 76 | 3 | 0.75 | 0.04003 |
| **GO:0032508** | AT1G06670, AT3G12810, AT5G45400 | DNA duplex unwinding | 76 | 3 | 0.75 | 0.04003 |
| **GO:2000113** | **AT2G19910, AT2G22670, AT2G25900, AT3G07100, AT3G12280, AT3G61050, AT5G22650, AT5G22220** | **negative regulation of cellular macromol...** | **388** | **8** | **3.85** | **0.04065** |
| **GO:0140014** | AT1G15660, AT3G16730, AT3G55340, AT5G10010, AT5G15540, AT5G37630 | mitotic nuclear division | 142 | 6 | 1.41 | 0.04083 |
| **GO:0010605** | **AT2G02090, AT2G19910, AT2G22670, AT2G25900, AT2G32710, AT2G46510, AT3G07100, AT3G12280, AT3G12810, AT3G61050, AT5G10010, AT5G22650, AT5G22220** | **negative regulation of macromolecule met...** | **765** | **13** | **7.6** | **0.04268** |
| **GO:0090698** | **AT1G80080, AT2G22670, AT3G12280, AT3G22780, AT5G16780** | **post-embryonic plant morphogenesis** | **192** | **5** | **1.91** | **0.04307** |
| **GO:0009414** | **AT1G69600, AT2G18960, AT2G39800, AT2G46680, AT3G26090, AT3G61050, AT5G08620, AT5G60410** | **response to water deprivation** | **393** | **8** | **3.9** | **0.04327** |
| **GO:0048583** | **AT1G60995, AT1G80080, AT2G39940, AT3G16730, AT3G17609, AT3G26090, AT3G55150, AT4G13040, AT5G10010, AT5G35580, AT5G35980, AT5G37190, AT5G60410** | **regulation of response to stimulus** | **768** | **13** | **7.63** | **0.04379** |
| **GO:0015931** | **AT1G06890, AT2G05120, AT2G33750, AT3G10650, AT5G51200** | **nucleobase-containing compound transport** | **193** | **5** | **1.92** | **0.04387** |
| **GO:0044275** | AT1G14520, AT1G69830, AT4G39010 | cellular carbohydrate catabolic process | 80 | 3 | 0.79 | 0.04547 |
| **GO:0034654** | **AT5G57160, AT2G22670, AT3G06380, AT4G23160, AT5G04670, AT3G16980, AT3G31430, ATMG00860, ATMG00300, AT1G49900, AT5G37190, AT5G42905, AT4G16845, AT1G06740, AT5G22650, AT5G15690, AT2G46680, AT1G48720, AT4G13040, AT3G09820, AT4G00980, AT4G10070, AT4G08540, AT5G24120, AT4G29090, AT2G05120, AT1G64490, AT3G22780, AT2G25900, AT2G19910, AT5G11430, AT5G64420, AT1G55325, AT3G61050, AT3G52200, AT5G22220, AT3G12270, AT3G12280, AT1G69600, AT4G04650, AT2G46510, AT5G58850, AT5G45400, AT5G15540, AT5G35170, AT2G01370, AT3G17609** | **nucleobase-containing compound biosynthe...** | **3106** | **47** | **30.84** | **0.04680** |
| **GO:0009415** | **AT1G69600, AT2G18960, AT2G39800, AT2G46680, AT3G26090, AT3G61050, AT5G08620, AT5G60410** | **response to water** | **401** | **8** | **3.98** | **0.04768** |

| | | | | | | |
|---|---|---|---|---|---|---|
| **GO:0018130** | **AT5G57160, AT2G22670, AT3G06380, AT4G23160, AT5G04670, AT3G16990, AT3G16980, AT3G31430, ATMG00860, ATMG00300, AT1G49900, AT5G37190, AT5G14800, AT5G42905, AT4G16845, AT1G06740, AT5G22650, AT5G15690, AT2G46680, AT1G48720, AT4G13040, AT3G09820, AT4G00980, AT4G10070, AT4G08540, AT5G24120, AT4G29090, AT2G39800, AT2G05120, AT1G64490, AT3G22780, AT2G25900, AT2G19910, AT5G11430, AT5G64420, AT1G55325, AT3G61050, AT3G52200, AT5G22220, AT3G12270, AT3G12280, AT1G69600, AT4G04650, AT2G46510, AT2G46580, AT5G58850, AT5G45400, AT5G15540, AT5G35170, AT2G01370, AT3G17609** | **heterocycle biosynthetic process** | **3334** | **51** | **33.11** | **0.04771** |
| **GO:0010558** | **AT2G22670, AT2G19910, AT2G25900, AT3G07100, AT3G12280, AT3G61050, AT5G22650, AT5G22220** | **negative regulation of macromolecule bio...** | **403** | **8** | **4** | **0.04882** |
| **GO:0006302** | AT4G05360, AT5G15540, AT5G45400, AT5G57160 | double-strand break repair | 138 | 4 | 1.37 | 0.04904 |
| **GO:0030641** | AT2G01980, AT2G18960 | regulation of cellular pH | 36 | 2 | 0.36 | 0.04960 |
| **GO:0051453** | AT2G01980, AT2G18960 | regulation of intracellular pH | 36 | 2 | 0.36 | 0.04960 |
| **GO:0006081** | AT2G46580, AT3G16910, AT4G14850 | cellular aldehyde metabolic process | 83 | 3 | 0.82 | 0.04978 |

**Table S5.** Phenotype scores for *Cochlearia* plants following salt stress treatment.

| Maternal Line | Scores | Average Score | Number of Samples | Standard Deviation | Ploidy |
|---|---|---|---|---|---|
| CHA_001 | 7, 5, 6, 7, 5, 7, 7, 4, 4, 3, 5, 8, 0, 0, 3 | 4.73 | 15 | 2.46 | 2x |
| CHA_010 | 5, 7, 5, 6, 4, 2, 3 | 4.57 | 7 | 1.72 | 2x |
| CHA_011 | 7, 4, 0, 4, 6, 6, 8, 7, 2, 2, 4, 5, 3 | 4.46 | 13 | 2.33 | 2x |
| CHA_002 | 4, 6, 6, 7, 5, 5, 6, 6 | 5.63 | 8 | 0.92 | 2x |
| CHA_008 | 7, 7, 8, 8, 8, 8, 7, 7, 9, 9, 7, 7, 9, 6, 9, 8, 6, 8, 8 | 7.68 | 19 | 0.95 | 2x |
| CWY_001 | 1 | 1.00 | 1 | N/A | 4x |
| CWY_013 | 3, 3, 3, 2, 2, 3, 2, 2, 5, 5, 5 | 3.18 | 11 | 1.25 | 4x |
| CWY_002 | 2 | 2.00 | 1 | N/A | 4x |
| CWY_003 | 3, 4, 5, 5, 2, 4, 3, 4, 4, 2, 3, 2, 1 | 3.23 | 13 | 1.24 | 4x |
| CWY_005 | 4, 3, 1, 2, 3, 2, 2, 3, 1, 1, 1 | 2.09 | 11 | 1.04 | 4x |
| LAB_002 | 5, 7, 7 | 6.33 | 3 | 1.15 | 2x |
| LAB_003 | 6, 4, 8, 6, 5, 5, 6, 8, 6, 5, 5, 4, 6, 2, 7, 4, 5, 2, 4, 7 | 5.25 | 20 | 1.65 | 2x |
| LAB_007 | 5, 6, 5, 7, 4, 6, 3, 6, 5, 3, 4, 4, 5, 5, 6, 6 | 5.00 | 16 | 1.15 | 2x |
| LAB_008 | 8, 6, 6, 7, 7, 7, 8, 8, 5, 7, 6, 8, 7, 6, 7, 7, 5, 7, 7, 7 | 6.80 | 20 | 0.89 | 2x |
| LAB_009 | 6, 5, 4, 5, 5, 5, 6, 7, 8, 5, 6, 5, 6, 5, 4, 5 | 5.44 | 16 | 1.03 | 2x |
| LAM_017 | 8, 6 | 7.00 | 2 | 1.41 | 2x |
| MAL_002 | 6, 7, 5, 6, 6, 7, 8, 6, 7, 5, 2, 0, 4, 6, 6, 7 | 5.50 | 16 | 2.03 | 2x |
| MAL_003 | 7, 8, 8, 5, 5, 5, 8, 8, 8, 5, 6, 9, 5 | 6.69 | 13 | 1.55 | 2x |
| MAL_004 | 7, 6, 6, 6, 4, 4, 6, 7, 6, 7, 7 | 6.00 | 11 | 1.10 | 2x |
| MAL_007 | 6, 5, 7, 6, 8, 6, 3, 4, 6, 6, 4, 6, 1, 3, 5 | 5.07 | 15 | 1.79 | 2x |
| NEN_002 | 1, 3, 2, 2, 3, 3, 3, 3, 1, 2, 1, 1, 2, 3, 3, 2, 3, 1, 2, 1 | 2.10 | 20 | 0.85 | 2x |
| NEN_003 | 7, 8, 8, 6, 6, 6, 7, 5, 8, 5, 4, 6, 8, 7, 0, 3, 7, 8, 5, 6 | 6.00 | 20 | 2.00 | 2x |
| NEN_004 | 2, 5, 7, 5, 4, 6, 7, 5, 5, 8, 8, 6, 6 | 5.69 | 13 | 1.65 | 2x |
| NEN_005 | 9, 6, 3, 7, 4, 6, 5, 7, 5, 8, 9, 6, 7, 3, 0, 4 | 5.56 | 16 | 2.39 | 2x |
| NEN_006 | 4, 6, 5, 6 | 5.25 | 4 | 0.96 | 2x |
| SCU_012 | 4, 3, 5, 4, 6, 4, 5, 4, 5, 5, 6, 7, 5, 5, 5, 5, 0, 5, 4, 7 | 4.70 | 20 | 1.49 | 4x |
| SCU_020 | 6, 7, 7, 7, 7, 7, 9, 7, 8, 6, 9, 10, 8, 9, 8, 8, 7, 9, 9 | 7.79 | 19 | 1.13 | 4x |
| SCU_003 | 4, 4, 3, 1 | 3.00 | 4 | 1.41 | 4x |
| SCU_009 | 3, 2, 3, 5 | 3.25 | 4 | 1.26 | 4x |
| TMN_001 | 3, 3, 4, 6, 3, 3, 5, 5, 4, 3, 2, 3, 0, 2, 3, 3, 2, 3, 4 | 3.21 | 19 | 1.32 | 4x |
| TMN_003 | 4, 5, 5, 5, 3, 3, 3, 2, 6, 3, 3, 5, 4, 4, 5, 4, 5, 7, 5, 4 | 4.25 | 20 | 1.21 | 4x |
| TMN_005 | 1, 5, 2, 1 | 2.25 | 4 | 1.89 | 4x |
| TMN_006 | 5, 4 | 4.50 | 2 | 0.71 | 4x |
| TYN_0H1 | 1, 4, 8, 5, 4, 2, 2, 2, 3, 2, 2, 1, 1, 3, 1, 2, 1, 2, 1, 1 | 2.40 | 20 | 1.76 | 2x |
| TYN_0H4 | 2, 3, 1, 3, 2, 1, 3, 1, 0, 1, 2, 4, 2, 1, 3, 2, 1, 2, 2, 0 | 1.80 | 20 | 1.06 | 2x |
| TYN_0H5 | 1, 3, 9, 7, 9, 8, 4, 0, 7, 8, 8, 8, 7, 6, 8, 9, 5, 5, 6, 5 | 6.15 | 20 | 2.58 | 2x |
| TYN_0H7 | 0, 5, 6, 6, 7, 5, 5, 4, 3, 0, 4, 6, 3, 2, 4, 5, 4, 2, 5, 3 | 3.95 | 20 | 1.90 | 2x |
| TYN_0H8 | 7, 6, 3, 4, 3, 3, 4, 5, 6, 6, 4, 5, 4, 3, 0, 1, 6, 5, 6, 6 | 4.35 | 20 | 1.81 | 2x |
| All diploids | N/A | 4.98 | 367 | 1829.00 | 2x |
| All tetraploids | N/A | 4.14 | 129 | 534.00 | 4x |

**Table S6**. Net Photosynthetic rates under drought relative to control conditions.

| relative net Photosynthesis | | | | | | | |
|---|---|---|---|---|---|---|---|
| **Species** | **Treatment** | **Ploidy** | **N** | **Average** | **sd** | **se** | **ci** |
| ELI21 | Control | 4x | 4 | **1.00** | 0.12 | 0.06 | 0.19 |
| ELI21 | Drought | 4x | 5 | **0.91** | 0.10 | 0.04 | 0.12 |
| ELI23 | Control | 4x | 3 | **1.00** | 0.16 | 0.09 | 0.40 |
| ELI23 | Drought | 4x | 4 | **0.78** | 0.04 | 0.02 | 0.06 |
| ELI25 | Control | 4x | 5 | **1.00** | 0.22 | 0.10 | 0.27 |
| ELI25 | Drought | 4x | 3 | **0.99** | 0.09 | 0.05 | 0.22 |
| ERS21+22 | Control | 4x | 4 | **1.00** | 0.05 | 0.03 | 0.08 |
| ERS21+22 | Drought | 4x | 4 | **1.04** | 0.10 | 0.05 | 0.17 |
| ERS23 | Control | 4x | 3 | **1.00** | 0.19 | 0.11 | 0.48 |
| ERS23 | Drought | 4x | 4 | **0.66** | 0.04 | 0.02 | 0.07 |
| ERS26 | Control | 4x | 3 | **1.00** | 0.27 | 0.15 | 0.66 |
| ERS26 | Drought | 4x | 4 | **0.83** | 0.10 | 0.05 | 0.16 |
| LAB22 | Control | 2x | 5 | **1.00** | 0.25 | 0.11 | 0.31 |
| LAB22 | Drought | 2x | 5 | **0.77** | 0.11 | 0.05 | 0.14 |
| LAB23 | Control | 2x | 4 | **1.00** | 0.06 | 0.03 | 0.09 |
| LAB23 | Drought | 2x | 5 | **0.60** | 0.17 | 0.08 | 0.22 |
| Nent20 | Control | 2x | 3 | **1.00** | 0.03 | 0.02 | 0.08 |
| Nent20 | Drought | 2x | 7 | **0.54** | 0.14 | 0.05 | 0.13 |
| Nent23 | Control | 2x | 4 | **1.00** | 0.07 | 0.03 | 0.11 |
| Nent23 | Drought | 2x | 3 | **0.48** | 0.17 | 0.10 | 0.42 |
| Nent24 | Control | 2x | 4 | **1.00** | 0.32 | 0.16 | 0.51 |
| Nent24 | Drought | 2x | 6 | **0.86** | 0.12 | 0.05 | 0.12 |
| PIT_020 | Control | 2x | 13 | **1.00** | 0.14 | 0.04 | 0.09 |
| PIT_020 | Drought | 2x | 15 | **0.82** | 0.23 | 0.06 | 0.13 |
| PIT_022 | Control | 2x | 3 | **1.00** | 0.13 | 0.07 | 0.32 |
| PIT_022 | Drought | 2x | 3 | **0.58** | 0.08 | 0.05 | 0.20 |
| PIT_023 | Control | 2x | 3 | **1.00** | 0.03 | 0.02 | 0.08 |
| PIT_023 | Drought | 2x | 3 | **0.84** | 0.07 | 0.04 | 0.19 |
| PIT_025 | Control | 2x | 5 | **1.00** | 0.07 | 0.03 | 0.08 |
| PIT_025 | Drought | 2x | 5 | **0.62** | 0.04 | 0.02 | 0.05 |
| SKN20 | Control | 4x | 3 | **1.00** | 0.18 | 0.10 | 0.44 |
| SKN20 | Drought | 4x | 6 | **0.65** | 0.10 | 0.04 | 0.10 |
| SKN21-A | Control | 4x | 6 | **1.00** | 0.16 | 0.07 | 0.17 |
| SKN21-A | Drought | 4x | 6 | **0.87** | 0.20 | 0.08 | 0.21 |
| SKN21-B | Control | 4x | 3 | **1.00** | 0.14 | 0.08 | 0.34 |
| SKN21-B | Drought | 4x | 5 | **0.93** | 0.27 | 0.12 | 0.34 |
| SKN24 | Control | 4x | 5 | **1.00** | 0.12 | 0.06 | 0.15 |
| SKN24 | Drought | 4x | 5 | **0.82** | 0.16 | 0.07 | 0.20 |

**Note:** Averages, standard deviation and 95% confidence interval under control and drought stress are shown.

**Table S7.** Stomatal conductance under drought relative to control conditions.

| relative Stomata Conductance | | | | | | | | |
|---|---|---|---|---|---|---|---|---|
| **Species** | **Treatment** | **Ploidy** | **N** | **Average** | **sd** | **se** | **ci** | **rn** |
| ELI21 | Control | 4x | 4 | **1.00** | 0.15 | 0.08 | 0.24 | 1.00 |
| ELI21 | Drought | 4x | 5 | **0.70** | 0.20 | 0.09 | 0.24 | 0.70 |
| ELI23 | Control | 4x | 3 | **1.00** | 0.17 | 0.10 | 0.42 | 1.00 |
| ELI23 | Drought | 4x | 4 | **0.98** | 0.17 | 0.09 | 0.28 | 0.98 |
| ELI25 | Control | 4x | 5 | **1.00** | 0.20 | 0.09 | 0.24 | 1.00 |
| ELI25 | Drought | 4x | 3 | **1.06** | 0.95 | 0.55 | 2.37 | 1.06 |
| ERS21+22 | Control | 4x | 4 | **1.00** | 0.31 | 0.15 | 0.49 | 1.00 |
| ERS21+22 | Drought | 4x | 4 | **0.62** | 0.11 | 0.05 | 0.17 | 0.62 |
| ERS23 | Control | 4x | 3 | **1.00** | 0.39 | 0.22 | 0.97 | 1.00 |
| ERS23 | Drought | 4x | 4 | **1.01** | 0.50 | 0.25 | 0.79 | 1.01 |
| ERS26 | Control | 4x | 3 | **1.00** | 0.25 | 0.14 | 0.61 | 1.00 |
| ERS26 | Drought | 4x | 4 | **1.44** | 0.82 | 0.41 | 1.31 | 1.44 |
| LAB22 | Control | 2x | 5 | **1.00** | 0.48 | 0.21 | 0.60 | 1.00 |
| LAB22 | Drought | 2x | 5 | **0.53** | 0.24 | 0.11 | 0.30 | 0.53 |
| LAB23 | Control | 2x | 4 | **1.00** | 0.41 | 0.20 | 0.65 | 1.00 |
| LAB23 | Drought | 2x | 5 | **0.39** | 0.09 | 0.04 | 0.12 | 0.39 |
| Nent20 | Control | 2x | 3 | **1.00** | 0.00 | 0.00 | 0.01 | 1.00 |
| Nent20 | Drought | 2x | 7 | **0.52** | 0.18 | 0.07 | 0.16 | 0.52 |
| Nent23 | Control | 2x | 4 | **1.00** | 0.17 | 0.08 | 0.26 | 1.00 |
| Nent23 | Drought | 2x | 3 | **0.54** | 0.18 | 0.11 | 0.46 | 0.54 |
| Nent24 | Control | 2x | 4 | **1.00** | 0.27 | 0.13 | 0.43 | 1.00 |
| Nent24 | Drought | 2x | 6 | **0.71** | 0.18 | 0.07 | 0.19 | 0.71 |
| PIT_020 | Control | 2x | 13 | **1.00** | 0.23 | 0.06 | 0.14 | 1.00 |
| PIT_020 | Drought | 2x | 15 | **0.81** | 0.26 | 0.07 | 0.14 | 0.81 |
| PIT_022 | Control | 2x | 3 | **1.00** | 0.43 | 0.25 | 1.07 | 1.00 |
| PIT_022 | Drought | 2x | 3 | **1.18** | 0.05 | 0.03 | 0.12 | 1.18 |
| PIT_023 | Control | 2x | 3 | **1.00** | 0.37 | 0.21 | 0.92 | 1.00 |
| PIT_023 | Drought | 2x | 3 | **0.71** | 0.38 | 0.22 | 0.95 | 0.71 |
| PIT_025 | Control | 2x | 5 | **1.00** | 0.15 | 0.07 | 0.19 | 1.00 |
| PIT_025 | Drought | 2x | 5 | **0.28** | 0.04 | 0.02 | 0.05 | 0.28 |
| SKN20 | Control | 4x | 3 | **1.00** | 0.31 | 0.18 | 0.78 | 1.00 |
| SKN20 | Drought | 4x | 6 | **0.70** | 0.08 | 0.03 | 0.08 | 0.70 |
| SKN21-A | Control | 4x | 6 | **1.00** | 0.38 | 0.15 | 0.40 | 1.00 |
| SKN21-A | Drought | 4x | 6 | **0.84** | 0.36 | 0.15 | 0.37 | 0.84 |
| SKN21-B | Control | 4x | 3 | **1.00** | 0.16 | 0.09 | 0.39 | 1.00 |
| SKN21-B | Drought | 4x | 5 | **0.91** | 0.33 | 0.15 | 0.41 | 0.91 |
| SKN24 | Control | 4x | 5 | **1.00** | 0.12 | 0.05 | 0.14 | 1.00 |
| SKN24 | Drought | 4x | 5 | **0.57** | 0.13 | 0.06 | 0.16 | 0.57 |

**Note:** Averages, standard deviation and 95% confidence intervals under control and drought stress are shown.

**Table S8.** Stomatal conductance and photosynthesis model results.

| | **rel. Stomatal Conductance** | | | **rel. Photosynthesis** | | |
|---|---|---|---|---|---|---|
| *Predictors* | *Estimates* | *CI* | *p* | *Estimates* | *CI* | *p* |
| (Intercept) | -0.29 | -0.55 – -0.03 | **0.030** | -0.31 | -0.57 – -0.05 | **0.021** |
| Ploidy [4x] | 0.62 | 0.24 – 1.00 | **0.002** | 0.66 | 0.28 – 1.04 | **0.001** |
| Observations | 98 | | | 98 | | |
| $R^2$ / $R^2$ adjusted | 0.097 / 0.088 | | | 0.110 / 0.100 | | |

**Table S9. Orthologue Overlaps.** Orthologue overlaps between *A. arenosa, C. amara* and *Cochlearia*.

| Orthogroup | C. amara gene(s) | Cochlearia gene(s) | A. arenosa gene(s) | Gene names and descriptions |
|---|---|---|---|---|
| **OG0000042** | CAg10694, CAg12636, CAg18129, CAg21234, CAg22036, CAg23713, CAg31403, CAg9211, CAg9746 | CPg12123, CPg123, CPg26718, **CPg30015**, CPg30849, CPg35203, CPg40035, CPg40038, CPg40041, CPg40044, CPg816 | AT1G80660.1, AT2G07560.1, **AT2G18960.1**, AT2G24520.1, AT3G42640.1, AT3G47950.1, AT4G30190.2, **AT5G57350.1**, AT5G62670.1 | **CPg30015, AT2G18960.1.** AHA1, H(+)-ATPASE 1, HA1, OPEN STOMATA 2, OST2, PLASMA MEMBRANE PROTON ATPASE, PMA. Encodes a plasma membrane proton ATPase. Mutants have a reduced ability to close their stomata in response to drought and are affected in stomatal but not seed responsiveness to ABA. The mRNA is cell-to-cell mobile. **AT5G57350.1.** AHA3, ARABIDOPSIS THALIANA ARABIDOPSIS H(+)-ATPASE, ATAHA3, H(+)-ATPASE 3, HA3. Member of Plasma membrane H+-ATPase family. |
| **OG0000137** | CAg10578, CAg15710, CAg28099 | CPg17784, **CPg20706**, CPg23114 | AT1G35375.1, **AT2G40955.1**, AT4G15096.1 | **CPg20706.** N/A. **AT2G40955.1.** Hypothetical protein. |
| **OG0001295** | CAg21238, CAg21241 | **CPg21082**, CPg26061 | AT3G47890.1, **AT3G47910.2** | **CPg21082, AT3G47890.2.** Ubiquitin carboxyl-terminal hydrolase-related protein. **AT3G47910.2.** Ubiquitin carboxyl-terminal hydrolase-related protein. |
| **OG0001840** | CAg11252, CAg7303 | **CPg1405**, CPg1515 | **AT3G16980.1**, AT4G16265.1 | **CPg1405, AT3G16980.1.** NRPB9A, NRPD9A, NRPE9A. One of two highly similar, non-catalytic subunits common to nuclear DNA-directed RNA polymerases II, IV and V; homologous to budding yeast RPB9. |
| **OG0002331** | CAg4288, **CAg5604** | CPg20737 | **AT5G08250.1**, AT5G23190.1 | **CAg5604, AT5G23190.1.** "CYTOCHROME P450, FAMILY 86, SUBFAMILY B, POLYPEPTIDE 1", CYP86B1. Cytochrome P450 CYP86B1. nuclear gene for chloroplast product. CYP86B1 is a very long chain fatty acid hydroxylase specifically involved in polyester monomer biosynthesis during the course of plant development. **AT5G08250.1.** Cytochrome P450 superfamily protein. |
| **OG0002450** | CAg34387, CAg34388 | CPg16666, **CPg18477**, CPg19816 | **AT4G10310.1** | **CPg18477, AT4G10310.1.** ATHKT1, HIGH-AFFINITY K+ TRANSPORTER 1, HKT1, HKT1;1. Encodes a sodium transporter (HKT1) expressed in xylem parenchyma cells. Mutants over-accumulate sodium in shoot tissue and have increased sodium in the xylem sap and reduced sodium in phloem sap and roots. |
| **OG0002593** | **CAg1480** | CPg35878, CPg4918 | **AT1G16460.2**, AT1G79230.1 | **CAg1480, AT1G16460.2.** ARABIDOPSIS THALIANA MERCAPTOPYRUVATE SULFURTRANSFERASE 2, ATMST2, ATRDH2, MERCAPTOPYRUVATE SULFURTRANSFERASE 2, MST2, RDH2, RHODANESE HOMOLOGUE 2, ST2, STR2, SULFURTRANSFERASE 2. Encodes a cytoplasmic thiosulfate:cyanide sulfurtransferase, activity of which increased the rhodanese activity of transgenic yeast. Can also act as a mercaptopyruvate sulfurtransferase. |
| **OG0003174** | **CAg20098** | CPg23931 | **AT3G59180.1**, AT3G59190.1 | **CAg20098, AT3G59190.1.** F-box/RNI-like superfamily protein. **AT3G59180.1.** Protein with RNI-like/FBD-like domain. |
| **OG0003528** | CAg4982 | **CPg21087** | **AT5G16030.3** | **CPg21087, AT5G16030.3.** Mental retardation GTPase activating protein. |
| **OG0004224** | CAg19850 | **CPg21389** | **AT3G61730.1**, AT5G36000.1 | **CPg21389, AT5G36000.1.** F-box protein RMF. **AT3G61730.1.** REDUCED MALE FERTILITY, RMF. Encodes a nuclear localized F-box protein that is involved in tapetal layer degeneration and pollen development. Interacts with ASK1 and that interaction is mediated by the F-box domain. |
| **OG0009635** | CAg18864 | **CPg18522** | **AT4G11110.1** | **CPg18522, AT4G11110.1.** SPA1-RELATED 2, SPA2. Encodes a member of the SPA (suppressor of phyA-105) protein family (SPA1-SPA4). SPA proteins contain an N-terminal serine/threonine kinase-like motif followed by a coiled-coil structure and a C-terminal WD-repeat domain. SPA proteins function redundantly in suppressing photomorphogenesis in dark- and light-grown seedlings. SPA2 primarily regulates seedling development in darkness and has little function in light-grown seedlings or adult plants. |
| **OG0010077** | **CAg4024** | CPg14116 | **AT5G05480.1** | **CAg4024, AT5G05480.1.** Peptide-N4-(N-acetyl-beta-glucosaminyl)asparagine amidase A protein. |

| | | | | |
|---|---|---|---|---|
| **OG0010187** | **CAg5641** | CPg21857 | **AT5G23570.1** | **CAg5641, AT5G23570.1.** ATSGS3, SGS3, SUPPRESSOR OF GENE SILENCING 3. Required for posttranscriptional gene silencing and natural virus resistance.SGS3 is a member of an 'unknown' protein family. Members of this family have predicted coiled coiled domains suggesting oligomerization and a potential zinc finger domain. Involved in the production of trans-acting siRNAs, through direct or indirect stabilization of cleavage fragments of the primary ta-siRNA transcript. Acts before RDR6 in this pathway. The mRNA is cell-to-cell mobile. |
| **OG0013525** | CAg7304 | **CPg1404** | **AT3G16990.1** | **CPg1404, AT3G16990.1.** Haem oxygenase-like, multi-helical. |
| **OG0015410** | CAg9024 | **CPg26954** | **AT5G64420.1** | **CPg26954, AT5G64420.1.** DNA polymerase V family. |
| **OG0015828** | CAg9293 | **CPg24947** | **AT5G61865.1** | **CPg24947, AT5G61865.1.** Hypothetical protein. |
| **OG0017644** | **CAg1401** | | **AT1G15580.1** | **CAg1401, AT1G69180.1.** CRABS CLAW, CRC. Putative transcription factor with zinc finger and helix-loop-helix domains, the later similar to HMG boxes. Involved in specifying abaxial cell fate in the carpel. Four putative LFY binding sites (CCANTG) and two potential binding sites for MADS box proteins known as CArG boxes (CC(A/T)6GG) were found in the region spanning 3.8 Kb upstream of the CRC coding region. CRC targets YABBY genes such as YUC4 in gynoecium development. |

**Note:** Genes in bold are 1% Fst candidate genes in the selection scans.

### Supplementary datasets

**Dataset S1.** Ploidy Survey.xlsx
**Dataset S2.** Cochlearia Fst Window Genes.xlsx
**Dataset S3.** Cochlearia MAV 1percent SNPs.xlsx
**Dataset S4.** *A_arenosa* Fst Window genes.xlsx
**Dataset S5.** *C_amara* Fst Window genes.xlsx
